# The inflammasome components NLRP3 and ASC act in concert with IRGM to rearrange the Golgi during viral infections

**DOI:** 10.1101/2020.04.27.063644

**Authors:** Coralie F. Daussy, Sarah C. Monard, Coralie Guy, Sara Muñoz-González, Maxime Chazal, Marit W. Anthonsen, Nolwenn Jouvenet, Thomas Henry, Marlène Dreux, Eliane F. Meurs, Marianne Doré Hansen

## Abstract

Hepatitis C virus (HCV) infection triggers Golgi fragmentation through the Golgi-resident protein immunity-related GTPase M (IRGM). Here, we report the role of NLRP3 and ASC, two inflammasome components, in the initial events leading to this fragmentation. We show that ASC resides at the Golgi with IRGM at homeostasis. Upon infection, ASC dissociates from both IRGM and Golgi and associates with HCV-induced NLRP3. NLRP3 silencing inhibits Golgi fragmentation. ASC silencing disrupts the Golgi structure in both control and infected cells and reduces the localization of IRGM at the Golgi. Silencing IRGM cannot totally restore the Golgi structure. These data highlight a role for ASC, upstream of the formation of the inflammasome, in regulating IRGM through its control on the Golgi. A similar mechanism occurs in response to Nigericin or infection with Zika virus (ZIKV). We propose a model for a newly ascribed function of the inflammasome components in Golgi structural remodeling.

## Introduction

Hepatitis C virus (HCV) is a major causative agent of chronic hepatitis and hepatocellular carcinoma worldwide. Twenty per cent of HCV-infected individuals can eliminate this virus but chronic infection occurs in 80% of cases. In the absence of efficient treatment such as the direct antiviral agents (DAA) therapy (Spengler, 2018), there is progressive degradation of the liver through inflammation, fibrosis, cirrhosis and hepatocarcinoma (Goossens & Hoshida, 2015). Chronic HCV infection remains a major global health problem in decades to come. HCV chronic infection is recognized to promote the dysregulation of lipid metabolism and thereby, can participate in many ways to the genesis of steatosis (Mirandola, Bowman, Hussain, & Alberti, 2010). Similar to all positive-strand RNA viruses studied thus far (Romero-Brey & Bartenschlager, 2014), HCV replicates its genome in association with intracellular membrane rearrangements. For HCV, the replication takes place within a so-called membranous web (MW) derived from the endoplasmic reticulum. The HCV MW has a complex morphology consisting of clusters of single-, double- and multi-membrane vesicles, which likely include autophagosomes, and is present at proximity to the lipid droplets (Nagy & Pogany, 2012; Romero-Brey et al., 2012). The MW formation is regulated by distinct HCV non-structural (NS) proteins through sequential interactions with several host factors (Nagy & Pogany, 2012; Reiss et al., 2011). For example, membrane-associated NS5A may bind directly to phosphatidylinositol-4 kinase III alpha (PI4KIIIα) to initiate morphogenesis of viral replication sites. Numerous other cellular components are subverted by HCV to promote MW formation. In addition, the HCV life cycle is also closely tied to lipid metabolism in infected cells (Lavie & Dubuisson, 2017).

We have recently reported that HCV exploits the immunity-related GTPase M (IRGM), previously identified as a risk factor for Crohn’s disease and tuberculosis, and for its role in autophagy (Chauhan, Mandell, & Deretic, 2016) to regulate the activity of the vesicular transport protein GBF1 and of the small Arf1 GTPase, thereby leading to Golgi fragmentation and facilitating HCV replication through lipid supply (Hansen et al., 2017). However, the initial events leading to the activation of IRGM upon HCV infection are not known.

Sensing of viral product or virus-induced cellular stress in cells of the immune system, such as monocytes and macrophages, can trigger the formation of the NLRP3 inflammasome, a complex of proteins involved in innate immune response leading to activation of caspase-1 and secretion of the pro-inflammatory cytokines IL-1ß and IL-18. Formation of the NLRP3 inflammasome complex was reported to occur also in hepatocytes, although of non-immune lineage, but in this case without secretion of the pro-inflammatory cytokines IL-1ß and IL-18 (Negash & Gale, 2015). In particular, HCV infection induces the activation of the NLRP3 inflammasome in cultured hepatoma cells that can ultimately lead to the caspase-1-mediated form of programmed cell death called pyroptosis (Kofahi, Taylor, Hirasawa, Grant, & Russell, 2016), Whatever the nature of the cells, activation of the NLRP3 inflammasome requires association of NLRP3 to caspase-1 through ASC (adaptor protein apoptosis-associated speck-like protein containing a CARD).

IRGM contains a globular N-terminal GTPase domain and, like ASC, a C-terminal helical Caspase Activation and Recruitment Domain (CARD). Here we show that IRGM interacts with ASC at the Golgi in homeostasis conditions. Upon HCV infection, NLRP3 is induced and ASC dissociates from both the Golgi and from IRGM. The ability of IRGM to mediate Golgi fragmentation was reduced in the absence of NLRP3 or ASC. Similar molecular events were observed in response to Nigericin, or upon infection of macrophages with Zika virus (ZIKV), thus assigning a newly defined function of the NLRP3 inflammasome components in the regulators of the host intracellular compartments upon infections.

## Results

### Golgi fragmentation upon HCV infection is dependent on NLRP3

Previous report showed that HCV activates the NLRP3 inflammasome in cultivated human hepatoma cells (Kofahi et al., 2016). Activation of the NLRP3 inflammasome requires first an increased expression of NLRP3, or priming, which can occur through NF-κB activation (H. Guo, Callaway, & Ting, 2015). Here, we have confirmed induction of NLRP3 by HCV in Huh7 cells, in association with augmented levels of HCV Core and HCV RNA (Fig 1A and B), and induction of IL8, used as a marker of NF-κB activation (Sun & Karin, 2008) (SFig 1A). Furthermore, we observed an induction of the interaction between NLRP3 and ASC upon HCV infection as measured by Proximity Ligation Assay (PLA) (Fig 1C, controls shown in SFig 2). Activation of caspase-1 in response to HCV infection was demonstrated using a fluorescent reporter of caspase-1 substrate (FAM-FLICA) and by comparison with a 3 hour-treatment with Nigericin, a well-known activator of the NLRP3 inflammasome (Mariathasan et al., 2006) (Fig 1D). Such a 3 hour-treatment with Nigericin was sufficient to trigger induction of IL8, chosen as a marker of NF-kB activation, and induce NLRP3 expression in Huh7 cells (SFig 1B, C). We have shown earlier that HCV triggers Golgi fragmentation to facilitate its replication through lipid supply (Hansen et al., 2017). To examine whether NLRP3 could play a role in Golgi fragmentation, HCV infection was performed in cells in which NLRP3 expression was silenced prior to infection. Reduced NLRP3 expression results in inhibition of HCV expression, as shown by immunoblot analysis of the HCV Core protein (Fig 1E). Furthermore, confocal analyses and quantification of Golgi phenotype revealed that the silencing of NLRP3 expression greatly impeded HCV-induced rearrangements of Golgi, as shown by decreased number of Golgi fragments, increased fragment area and decreased fragment circularity as compared to siCtrl-treated cells (Fig 1F). These results suggest the existence of a link between NLRP3 and the regulation of HCV induced-Golgi rearrangement.

**Fig 1.**
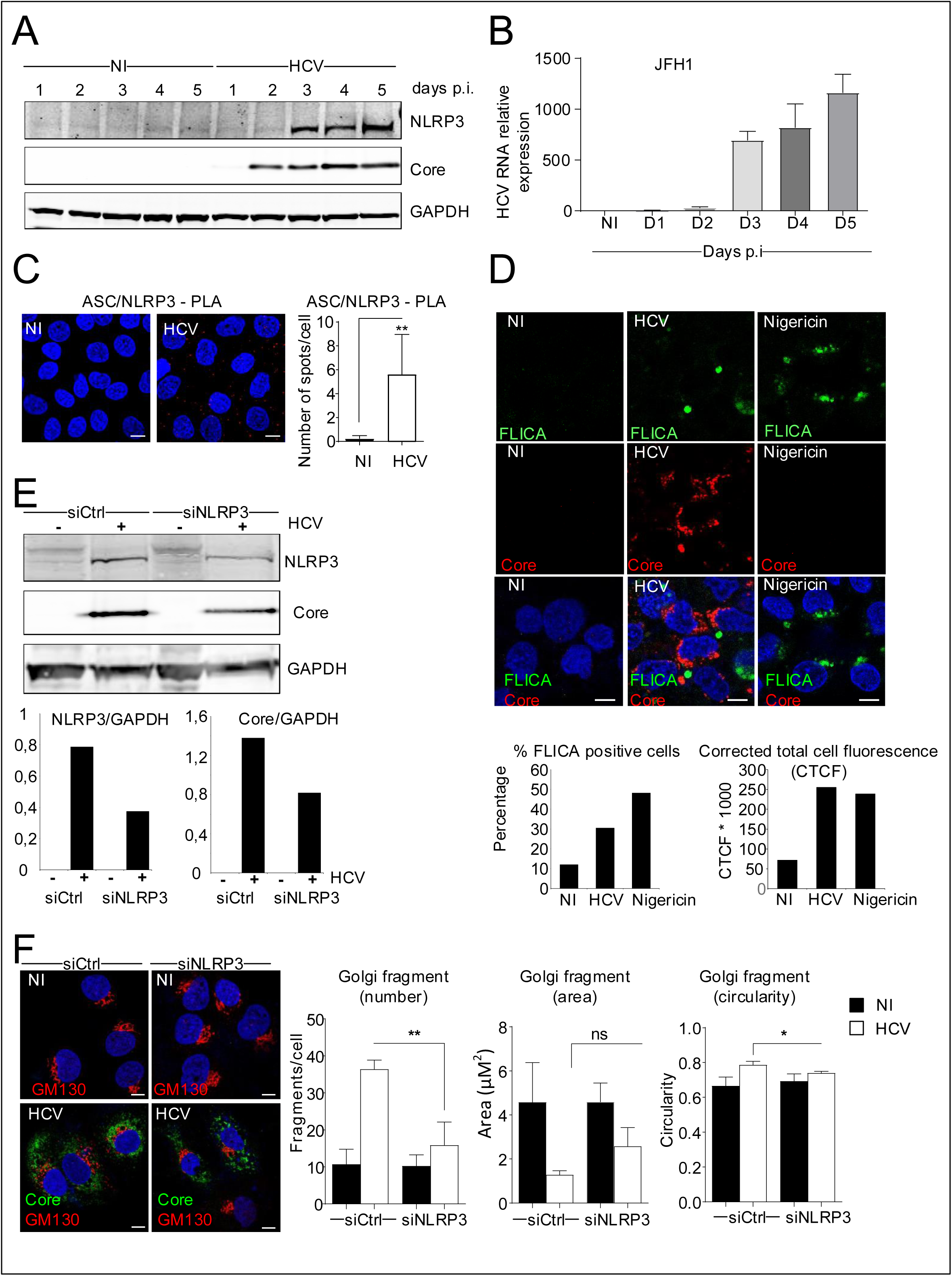
HCV infection activates the NLRP3 inflammasome and HCV-induced Golgi fragmentation is dependent on NLRP3. (A) The kinetics of expression of NLRP3 and HCV-Core protein in non-infected (NI) and HCV-infected Huh7 cells was examined by immunoblotting. GAPDH was used as a loading control. (B) HCV RNA levels were determined by RT-qPCR in cells infected with HCV. (C) Huh7 were infected with HCV for 5 days. Proximity Ligation Assay (PLA) was performed with antibodies directed against ASC and NLRP3, with the quantification of the number of PLA dots. Data shown are the mean ± SD, n=3 independent experiments, >180 cells per condition. (D) Huh7 cells were either infected with HCV (5 days) or treated with Nigericin (1 µM; 3 hours of treatment). Control cells were left uninfected and untreated. Cells were incubated for 1 hour with the FLICA reagent (stains the active form of caspase-1, green) prior to fixation. Fixed cells were immunostained with antibodies against HCV-Core protein (red) and nuclei stained with DAPI (blue). Percentage of infected cells positive for FLICA staining and the corrected total cell fluorescence (CTCF) for the FLICA staining was calculated. (E) Huh7 cells were reverse transfected with siRNAs against NLRP3 (siNLRP3) or control siRNA (siCtrl) prior to infection with HCV (MOI = 1) for 5 days. NLRP3 knockdown was controlled by immunoblotting and the effect of NLRP3 depletion on HCV protein levels was examined by immunoblotting with HCV-Core antibody. Protein levels were normalized to GAPDH. A blot from one representative experiment is shown. NLRP3 and HCV-Core levels were normalized to GAPDH (F) Representative images of cells depleted or not of NLRP3 and immunostained with antibodies against GM130 (red) and HCV-Core (green). DAPI staining marks nuclei (blue). The characteristics of Golgi fragments were calculated. Data shown are the means ± SD; n=3 independent experiments, >20 cells per condition.

**Fig 2.**
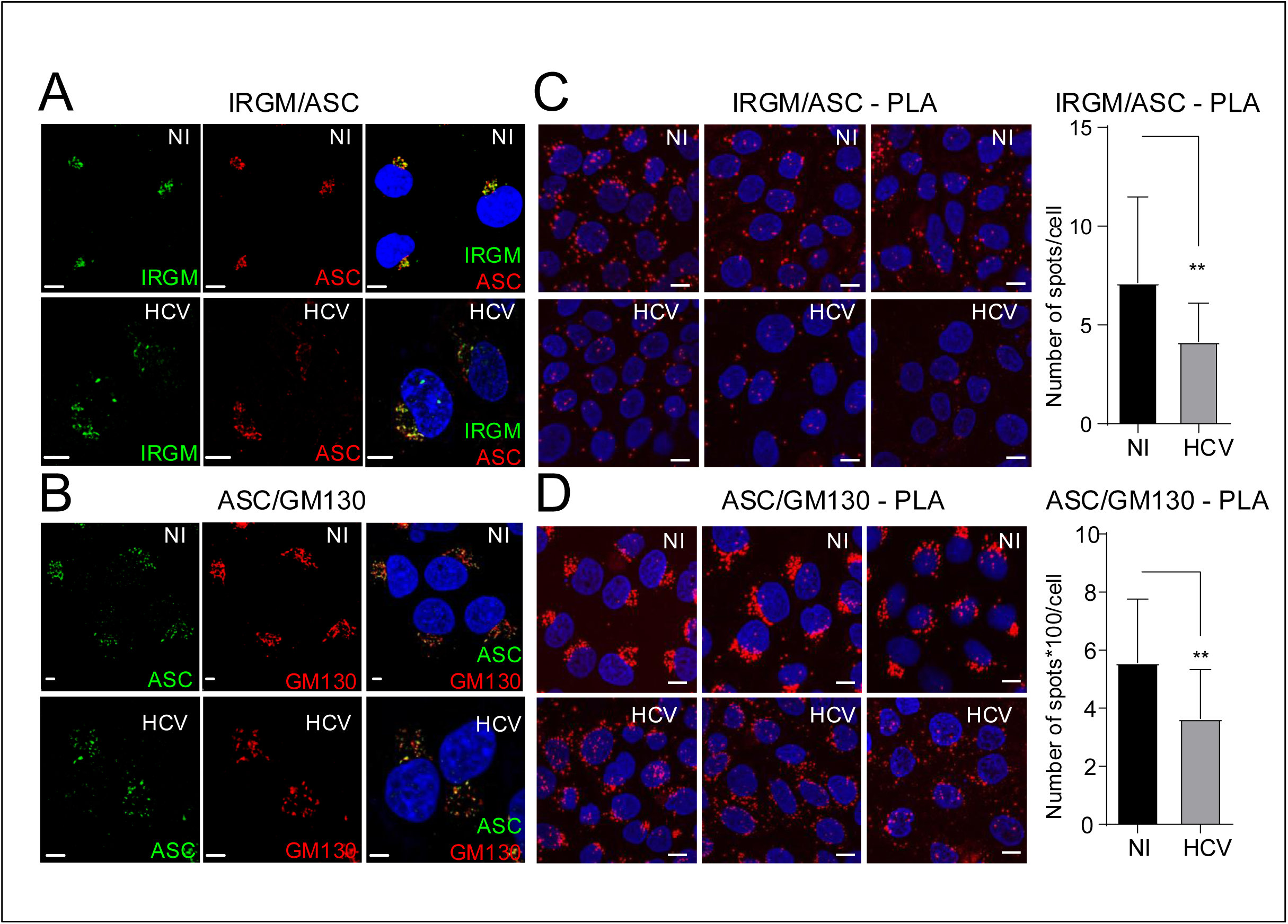
IRGM and ASC colocalize at the Golgi and separate upon HCV infection, together with Golgi fragmentation. (A, B) Huh7 cells were infected, or not, for 5 days. Representative images of cells immunostained with antibodies against IRGM (green) and ASC (red) or against ASC (green) and GM130 (red). DAPI staining marks nuclei (blue). (C, D) Proximity Ligation Assay (PLA) was performed with antibodies directed against ASC and IRGM or against ASC and GM130, with the quantification of the number of PLA dots. Data shown are the mean ± SD, n=3 independent experiments, >180 cells per condition. PLA performed on controls (cells untreated or incubated with each of the antibodies is shown on SFig 2)

### HCV infection triggers dissociation of ASC from IRGM at the Golgi

The formation of the NLRP3 inflammasome complex requires interaction of NLRP3 with ASC and caspase-1. Since both ASC and the immunity-related GTPase M (IRGM), which regulates the HCV-induced Golgi fragmentation (Hansen et al., 2017) contain a CARD domain, an interaction motif regulating several processes relating to innate responses, we examined the relationship between IRGM and ASC, during HCV infection using subcellular imaging analysis. Due to some uncertainties about the cellular localization of ASC from the literature (Misawa et al., 2013; Sagulenko et al., 2013; Zhou, Yazdi, Menu, & Tschopp, 2011), we first silenced its expression in the Huh7 cells (using three different siRNAs) before immunostaining. This resulted in a strong reduction of the expression of ASC both by immunofluorescence (SFig 3A) and by western blot (SFig 3B), thus confirming the specificity of our antibodies. We then observed colocalization between IRGM and ASC in the non-infected cells (Fig 2A). IRGM is a Golgi resident protein (Hansen et al., 2017), which might indicate that ASC can also localize at the Golgi. Localization of ASC at the Golgi was then confirmed by its strong colocalization with the cis-Golgi marker GM130 (Fig 2B). Interestingly, HCV infection impaired both the colocalization of ASC with IRGM (Fig 2A) and with the Golgi (Fig 2B). Quantification analysis of the IRGM/ASC association using PLA further showed that HCV infection significantly decreases the number of PLA spots per cell (Fig 2C). This revealed that HCV infection interferes with the IRGM/ASC association. Similarly, quantification analysis of the ASC/GM130 association showed a significant decrease of the frequency of ASC/GM130 PLA spots in the HCV-infected cells as compared to uninfected cells (Fig 2D). This shows that HCV infection disrupts, at least in part, the ASC/GM130 association. This stands in contrast to IRGM which remains associated with Golgi fragments upon HCV infection (Hansen et al., 2017). Altogether, these results reveal that ASC is a cis-Golgi resident protein, along with IRGM, and that HCV infection triggers its dissociation both from the Golgi apparatus and IRGM.

**Fig 3.**
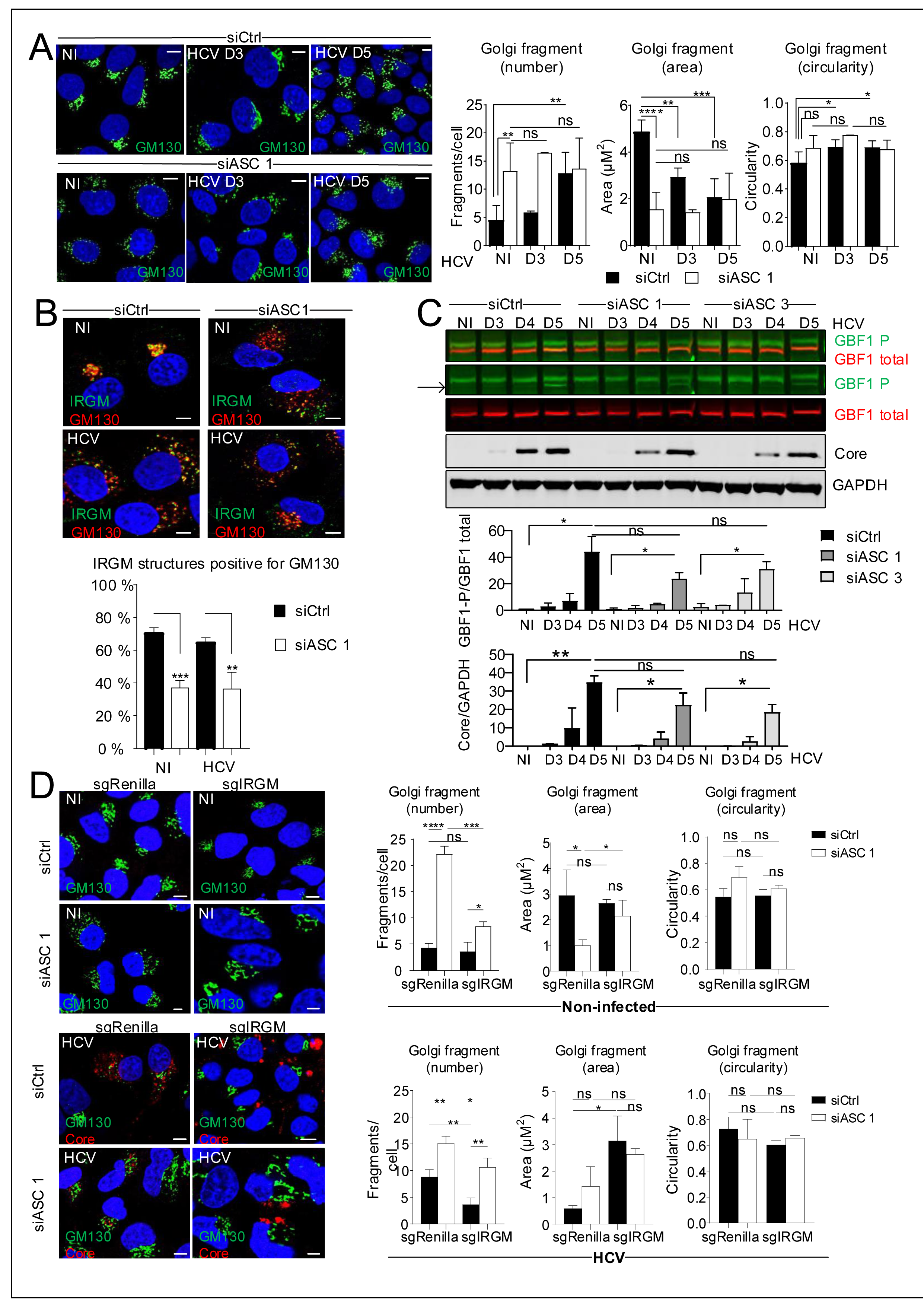
Role of ASC on Golgi fragmentation. Huh7 cells were reverse transfected with siRNAs against ASC (siASC) or control siRNA (siCtrl) prior to infection with HCV for 3 and 5 days. (A) Representative images of cells depleted or not of ASC and immunostained with antibodies against GM130 (green) and HCV-Core (not shown). DAPI staining marks nuclei (blue). The characteristics of Golgi fragments were calculated. Data shown are the means ± SD; n=5 independent experiments, >50 cells per condition. (B) The effect of siASC on GBF1 phosphorylation and on HCV protein levels were examined by immunoblotting. A merged image in the first row reveals overlap of signals (seen as orange) between phosphorylated (green) and total GBF1 (red) protein levels. The band corresponding to phospho-GBF1 (GBF1T1337) is indicated by an arrow. GBF1P1337 protein levels were normalized to total GBF1 protein levels. GAPDH was used as a loading control. (C) Representative images of cells depleted or not of ASC and immunostained with antibodies against IRGM (green) and GM130 (red). DAPI staining marks nuclei (blue). The MCC quantification method was used to quantify the number of IRGM structures positive for GM130. Data shown are the means ± SD; n=3 independent experiments, >20 cells per condition. (D) Representative images of CRISPR Huh7 sgIRGM and control sgRenilla cells depleted or not of ASC and infected or not with HCV for 3 days and immunostained with antibodies against GM130 (green) and HCV-Core (red). DAPI staining marks nuclei (blue). The characteristics of Golgi fragments were calculated. Data shown are the means ± SD; n=3 independent experiments, >20 cells per condition.

### ASC is involved in the control of the Golgi structure

We have shown that ASC associates with NLRP3 upon HCV infection and that the silencing of NLRP3 inhibits Golgi fragmentation in the HCV-infected cells (Fig 1F). We then examined the effect of silencing ASC on Golgi fragmentation. Huh7 cells were transfected with three different siRNA against ASC, and control siRNA as reference, prior to their infection or not with HCV for 3 and 5 days. While HCV infection triggered Golgi fragmentation in the siRNA Ctrl-treated cells (Fig 3A, SFig 4, black bars) as expected (Fig 1F and (Hansen et al., 2017)), the reduction of ASC expression led to a marked increase of Golgi fragmentation, both in the uninfected and HCV-infected cells, as demonstrated by higher proportion of small, circular Golgi fragments (Fig 3A, SFig 4). These data argue against a direct role of the NLRP3 inflammasome to trigger Golgi fragmentation and, instead, highlight an essential role for ASC in the control of the Golgi structure. This led us to question whether ASC could control the IRGM-induced Golgi fragmentation

**Fig 4.**
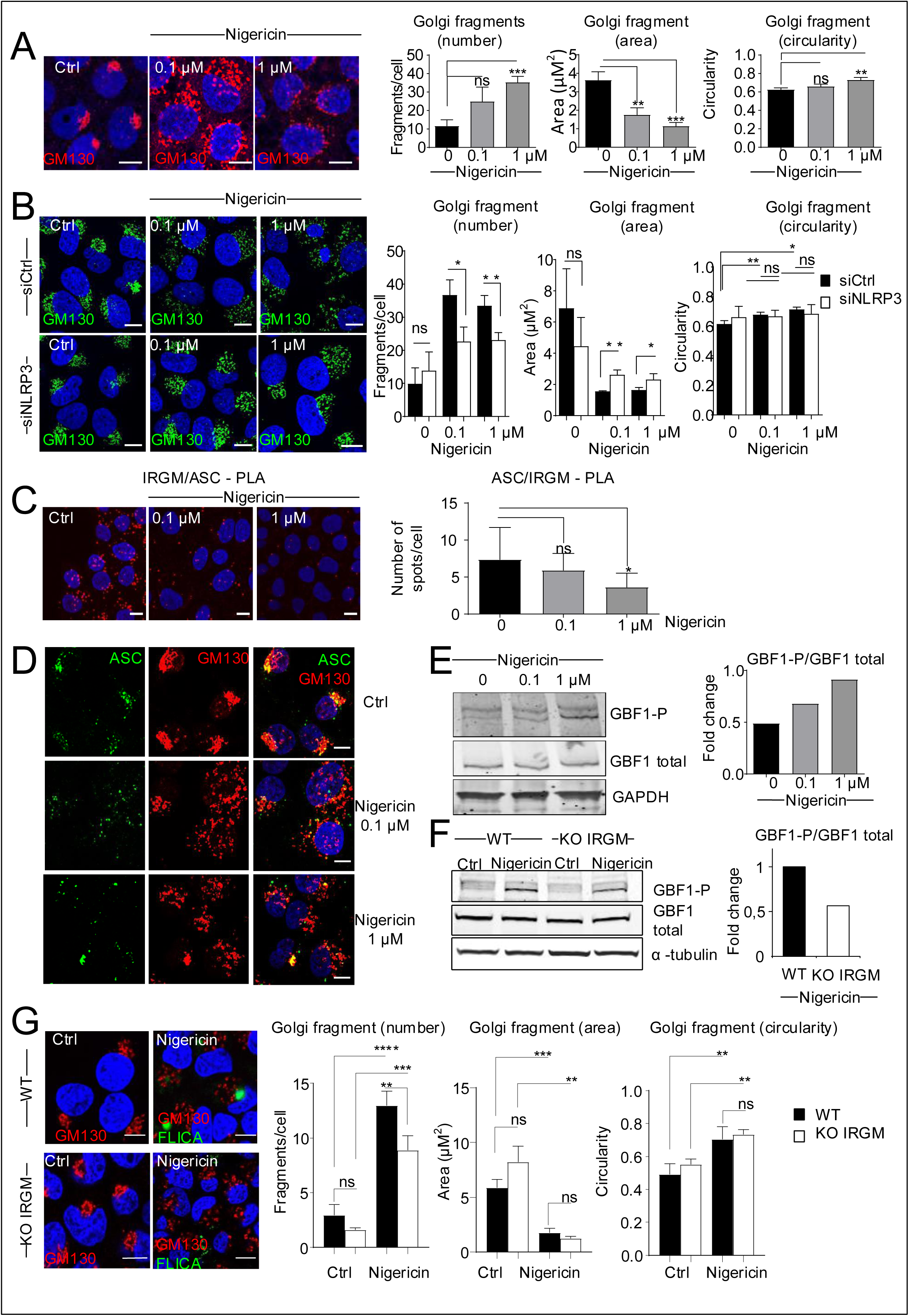
Golgi fragmentation upon Nigericin treatment of Huh7 cells is partly dependent on IRGM. (A) Representative images of cells non-treated (Ctrl) or treated with Nigericin (0.1 or 1 µM; 3 hours of treatment) and immunostained with an antibody against GM130 (red). DAPI staining marks nuclei (blue). The characteristics of Golgi fragments were calculated. Data shown are the means ± SD; n=3 independent experiments, >20 cells per condition. (B) Representative images of cells depleted or not of NLRP3 and non-treated (Ctrl) or treated with Nigericin (0.1 or 1 µM; 3 hours of treatment) was immunostained with antibodies against GM130 (green) and HCV-Core (not shown). DAPI staining marks nuclei (blue). The characteristics of Golgi fragments were calculated. Data shown are the means ± SD; n=3 independent experiments, >20 cells per condition. (C) Proximity Ligation Assay (PLA) was performed with antibodies directed against ASC and IRGM, with the quantification of the number of PLA dots. Data shown are the mean ± SD, n=3 independent experiments, >180 cells per condition. (D) Representative images of cells non-treated (Ctrl) or treated with Nigericin (0.1 or 1 µM; 3 hours of treatment) and immunostained with antibodies against ASC (green) and GM130 (red). DAPI staining marks nuclei (blue). (E) The effect of Nigericin on GBF1 phosphorylation was examined by immunoblotting using antibodies directed against the total and the phosphorylated (GBF1P1337) forms of GBF1. GAPDH was used as a loading control. GBF1P1337 protein levels were normalized to total GBF1 protein levels. (F) The effect of Nigericin (1 µM; 3 hours of treatment) on GBF1 phosphorylation was examined in CRISPR Huh7 sgIRGM and control sgRenilla cells by immunoblotting. Ctrl= Non-treated cells. α-tubulin was used as a loading control. GBF1P1337 protein levels were normalized to total GBF1 protein levels. (G) Representative images of CRISPR Huh7 sgIRGM and control sgRenilla cells treated or not with Nigericin (1 µM; 3 hours). Cells were incubated for 1 hour with the FAM-YVAD-FMK reagent (stains the active form of caspase-1, green) prior to fixation. Fixed cells were immunostained with GM130 (red). DAPI staining marks nuclei (blue). The characteristics of Golgi fragments were calculated. Data shown are the means ± SD; n=3 independent experiments, >20 cells per condition.

We examined the effect of silencing ASC on the localization of IRGM at the Golgi structure, in control cells and upon HCV infection. IRGM remains localized with the Golgi apparatus when intact or in HCV-induced Golgi fragments (Fig 3B; siCtrl), in agreement with our previous report (Hansen et al., 2017). In sharp contrast, ASC silencing greatly reduces the localization of IRGM at the Golgi in both the non-infected and HCV-infected cells (Fig 3B; siASC). This result suggests that ASC is pivotal for IRGM localization at the cis-Golgi (*i*.*e*. GM130 positive compartment).

We previously showed that the regulation of Golgi fragmentation by IRGM involved the phosphorylation of the vesicular transport protein GBF1, which is required to attract the Arf1 GTPase at the Golgi and to start the vesiculation process (Hansen et al., 2017). To examine the effect of ASC on this function of IRGM, Huh7 cells were silenced or not for ASC, submitted to kinetics of HCV infection and analyzed for GBF1 phosphorylation state and expression of HCV core. Our results showed that both GBF1 phosphorylation and expression of HCV Core protein levels were reduced in the absence of ASC as compared to the control cells (Fig 3C) but the difference was not found to be significant. These data, together with the 50% reduction of IRGM association with the Golgi in the absence of ASC, indicates that the presence of ASC is required for the stability of the association of IRGM with the Golgi but that it is not directly required for the ability of IRGM to mediate GBF1 phosphorylation and hence HCV expression.

Next, to further determine how ASC intersects with IRGM in the regulation of the Golgi structure, we examined the impact of the individual silencing of ASC *versus* combined with knocking-out (KO) IRGM, obtained by CRISPR–Cas9 approach. Consistently with our previous report (Hansen et al., 2017), IRGM KO prevents HCV-induced Golgi fragmentation (Fig 3D; black bars), and does not affect the Golgi structure in the control (WT) cells. Nonetheless, the silencing of ASC partially impedes the restoration of intact Golgi structure in IRGM KO cells and this was observed both in HCV-infected and non-infected cells (Fig 3D; white bars). Altogether, these data demonstrated a role for ASC in maintaining the Golgi structure at homeostasis (*i*.*e*. absence of infection) in part via its interaction and control of IRGM as well as via possible IRGM-independent regulation. Upon HCV infection, recruitment of ASC to NLRP3, may consequently lead to its dissociation from IRGM and favor the ability of IRGM to trigger GBF1 phosphorylation and formation of Golgi vesicles.

### Golgi fragmentation in response to Nigericin is mediated in part through the ASC/IRGM axis

Since we demonstrated that NLRP3, ASC and IRGM are key regulators for the Golgi shape in homeostasis or infection conditions, we then examined their regulation upon induction of the inflammasome by a virus-unrelated activation, using Nigericin (Mariathasan et al., 2006). Huh7 cells were submitted to Nigericin for 3 hours. Such time of treatment did not affect the cells morphology (not shown) and was sufficient to activate NF-κB, induce NLRP3 expression (SFig 1B, C) and activate the inflammasome (Fig 1D). The results showed a clear Golgi fragmentation in response to short-treatment by Nigericin with increased number of Golgi fragments and reciprocal decreased size and in a dose-dependent manner (Fig 4A). Next, confocal analyses and quantification of Golgi phenotype revealed that the silencing of NLRP3 expression significantly reduced the Nigericin-induced rearrangements of Golgi, as compared to siCtrl-treated cells (Fig 4B). These results suggest the existence of a link between NLRP3 and the regulation of Golgi rearrangement, similar to the situation with HCV infection. Then, we showed that Nigericin treatment triggers a significant dissociation of ASC from IRGM, as shown by PLA (Fig 4C) and also from the Golgi, as shown by confocal microscopy using an ASC antibody together with the cis-Golgi marker GM130 (Fig 4D). The ability of ASC proteins to dissociate from the Golgi upon Nigericin treatment was observed using other cell lines including macrophage derived from the monocytic U937 cell line (SFig 5), thus broadening our observations. Together these results further validate our hypothesis that displacement of ASC from Golgi is associated with Golgi fragmentation. We then showed that GBF1 phosphorylation was triggered upon Nigericin treatment in the Huh7 cells in a dose-dependent manner (Fig 4E) and was reduced in the IRGM-deficient cells (Fig 4F). Nonetheless, as opposed to HCV infection, the ability of Nigericin to trigger Golgi fragmentation was not totally inhibited by silencing IRGM (Fig 4G). Altogether, these data indicate that components of the inflammasome, such as NLRP3 and ASC, are involved, at least in part, in the process of the IRGM-Golgi fragmentation, in response to stress other than HCV infection.

**Fig 5.**
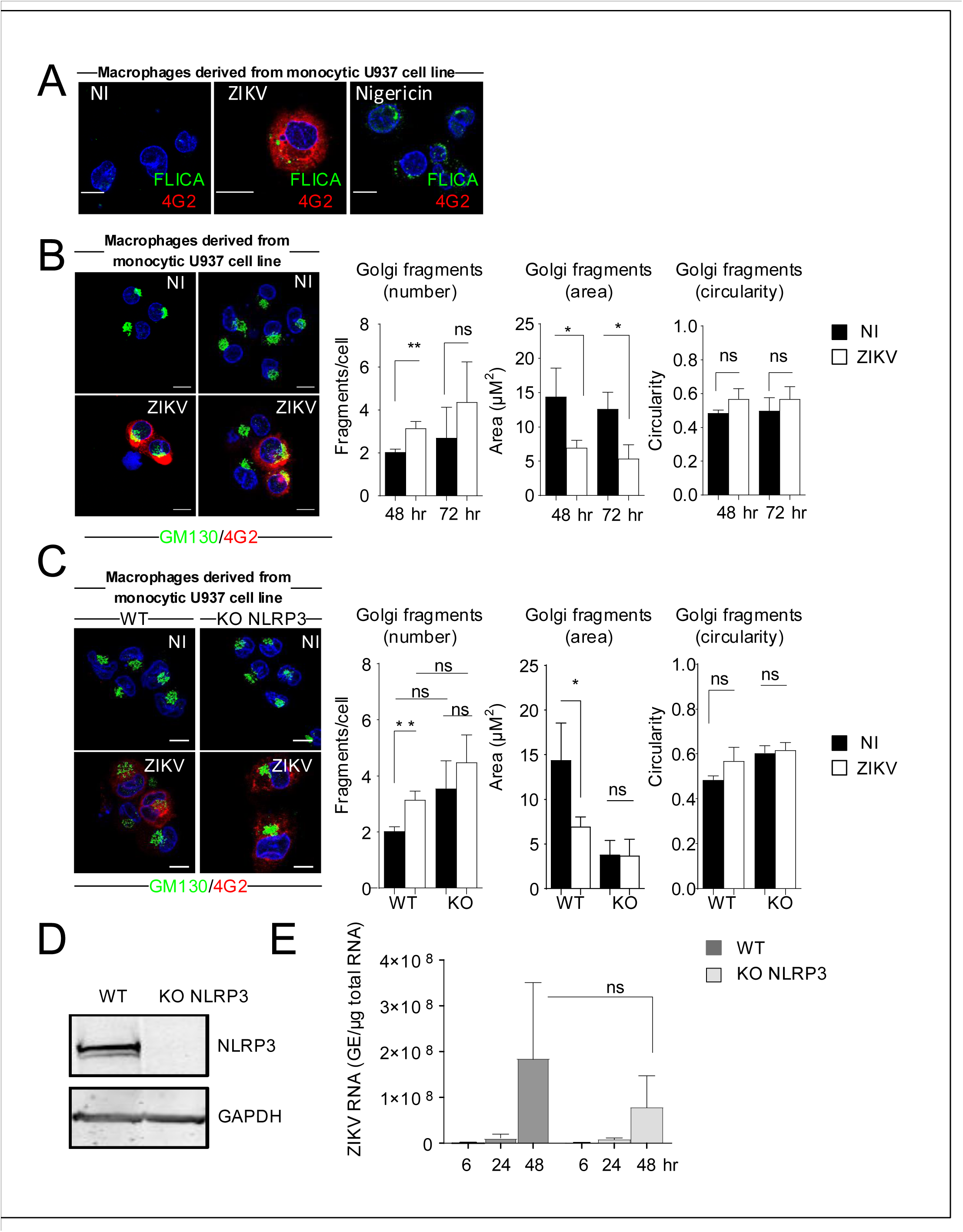
ZIKV infection triggers Golgi fragmentation through the NLRP3 inflammasome. (A) Macrophages derived from monocytic U937 cell line were either infected with ZIKV for 48hr or treated with Nigericin (1 µM; 3 hours of treatment). Control cells were left uninfected and untreated. Cells were incubated for 1 hour with the FAM-YVAD-FMK reagent (stains the active form of caspase-1, green) prior to fixation. Fixed cells were immunostained with antibodies against Flavivirus Envelope protein (4G2; red) and nuclei stained with DAPI (blue). (B) Representative images of macrophages derived from monocytic U937 cells infected for 48h or 72h with ZIKV and immunostained with antibodies against GM130 (green) and 4G2 (red). DAPI staining marks nuclei (blue). The characteristics of Golgi fragments were calculated. Data shown are the means ± SD; n=3 independent experiments, >20 cells per condition. (C) Representative images of macrophages derived from monocytic U937 cells KO for NLRP3 and parental cells immunostained with antibodies against GM130 (green) and 4G2 (red) after infection for 48h with ZIKV. DAPI staining marks nuclei (blue). The characteristics of Golgi fragments were calculated. Data shown are the means ± SD; n=3 independent experiments, >20 cells per condition. (D). NLRP3 knockdown was controlled by immunoblotting. GAPDH was used as a loading control. (E). ZIKV RNA levels were determined by RT-qPCR in the macrophages derived from monocytic U937 cells KO for NLRP3 and parental cells infected with ZIKV for the indicated duration.

### Golgi fragmentation through NLRP3 in macrophages in response to ZIKV infection

We then examined whether other viral infections can trigger Golgi fragmentation through NLRP3 and, in particular, whether this occurs in monocytic cell types, in which the inflammasome is well-studied. For this, we used ZIKA virus (ZIKV), another member of the *Flaviviridae* family, which notably infects monocytes/macrophages (Wang et al., 2018) thought to be pivotal for viral dissemination. ZIKV has been reported to trigger activation of the NLRP3 inflammasome and Golgi fragmentation (Sanchez-San Martin et al., 2018; Wang et al., 2018). Nonetheless, the relationship between the two events is still unknown. We confirmed that ZIKV infection activates the inflammasome (Fig 5A) and generates Golgi fragmentation in macrophages derived from the monocytic U937 cell line (Fig 5B). To evaluate the role of NLRP3 in the ZIKV-mediated Golgi fragmentation, ZIKV infection was carried out in U937 WT and KO NLRP3 (Rossignol, Peters, Connor, & Bullitt, 2017). Results showed that, upon ZIKV infection, Golgi structure was intact in these NLRP3 KO cells as compared to the WT counterparts (Fig 5C). The absence of NLRP3 expression in the macrophages derived from the monocytic U937 cell line KO NLRP3 was assessed by immunoblot (Fig 5D). This indicates that ZIKV infection can trigger Golgi fragmentation through NLRP3 in macrophages derived from monocytic U937 cell line. Viral replication was diminished but not significantly decreased by the reduced NLRP3 expression (Fig 5E). Altogether, these data indicate that Golgi fragmentation can occur through exploitation of inflammasome components in both non-hematopoietic and hematopoietic cells in response to different viral infections.

## Discussion

Here, we have demonstrated that HCV infection triggers the IRGM-mediated Golgi fragmentation through NLRP3 and ASC, two proteins which are usually known to be involved in the formation of inflammasome and we uncovered a role for ASC in the control of the Golgi structure. We showed that, while HCV infection provides all conditions to favor the formation of the NLRP3 inflammasome (induction of NLRP3, interaction NLRP3/ASC, FAM-FLICA labeling), only silencing of NLRP3 inhibits the HCV-mediated Golgi fragmentation, similar to the phenotype we described previously upon IRGM silencing (Hansen et al., 2017). Silencing of ASC did not inhibit Golgi fragmentation and, on the contrary, triggered this fragmentation, even in non-infected cells. This precludes a role for the inflammasome in the HCV-mediated Golgi fragmentation, since this would have been prevented, whether by silencing NLRP3 or ASC. We then showed that ASC and IRGM, which are two CARD-containing proteins, can physically interact and that they colocalize in homeostasis conditions at the Golgi apparatus. Previous studies on the intracellular localization of ASC were limited and somewhat contradictory, placing ASC in the cytosol, at the mitochondria or in the nucleus (Misawa et al., 2013; Sagulenko et al., 2013; Zhou et al., 2011). Using different siRNAs against ASC to assess the specificity of our anti ASC antibodies, we unambiguously showed that ASC resides at the Golgi in homeostatic conditions, in association with the cis-Golgi resident IRGM and GM130. HCV infection triggers an increased association between ASC and the HCV-induced NLRP3, the dissociation of ASC from the Golgi and from IRGM whilst IRGM still localizes to the Golgi. It is possible that dissociation of ASC from the Golgi and IRGM results from a change in the structure of the Golgi due to the presence of NLRP3 or directly because of the recruitment of ASC by NLRP3. The mechanisms leading to the formation of the NLRP3 inflammasome upon HCV infection are not known yet, but could be related to modifications of NLRP3, such as its phosphorylation, a step that is required for its deubiquitination to favor its oligomerization. NLRP3 is notably phosphorylated by JNK1 (Song et al., 2017). HCV infection can indeed trigger JNK activation (Chusri et al., 2016). NLRP3 can also be phosphorylated by protein kinase D. In this case, NLRP3 was found to be localized at MAMs, close to the Golgi, where PKD was itself recruited by diacylglycerol increase at the Golgi. Phosphorylated NLRP3 is then released and can be engaged in inflammasome (Zhang et al., 2017). Activation of PKD has been previously reported in HCV infection but was shown as inhibiting HCV secretion through the phosphorylation of OSBP (Amako, Syed, & Siddiqui, 2011). Here we have not addressed the phosphorylation change in NLRP3 upon HCV infection, but we show that its expression is induced upon HCV infection, that it associates with ASC and importantly that its depletion by silencing prevents Golgi fragmentation. Alternatively, the HCV NS3 protein has been reported to interact with IRGM (Gregoire et al., 2011) and also with GBF1 (Lebsir et al., 2019) and could play a role in the activation of the NLRP3 inflammasome. Whatever the case, since we have shown a direct link between NLRP3 and the IRGM-mediated Golgi fragmentation, our data emphasize the importance of NLRP3 in HCV infection and highlight the role of the ASC/IRGM interaction at the Golgi.

Upon HCV infection, dissociation of ASC from the Golgi may consequently lead to its dissociation from IRGM and favor the ability of IRGM to trigger GBF1 phosphorylation and formation of Golgi vesicles (Fig. 6). Since the intact Golgi structure was only partially restored by silencing IRGM in control cells silenced for ASC, this demonstrates that the role of ASC is not to directly regulate IRGM function through its association but rather to control the maintenance of the Golgi structure in absence of inflammasome activation. We hypothesize that in response to HCV infection, only a fraction of ASC population is released from IRGM (and probably from other partners) to associate with NLRP3 to form the inflammasome. Consequently, IRGM would remain associated to sufficiently organized Golgi structures to be able to mediate some Golgi fragmentation through GBF1 phosphorylation. We showed that NLRP3 was also involved in Golgi fragmentation upon treatment with Nigericin or infection with ZIKV. Altogether, our results highlight the key functional cross-talk between inflammasome components and Golgi fragmentation during viral infections in both non-hematopoeitic and hematopoeitic cells (Fig. 6).

**Figure 6.**
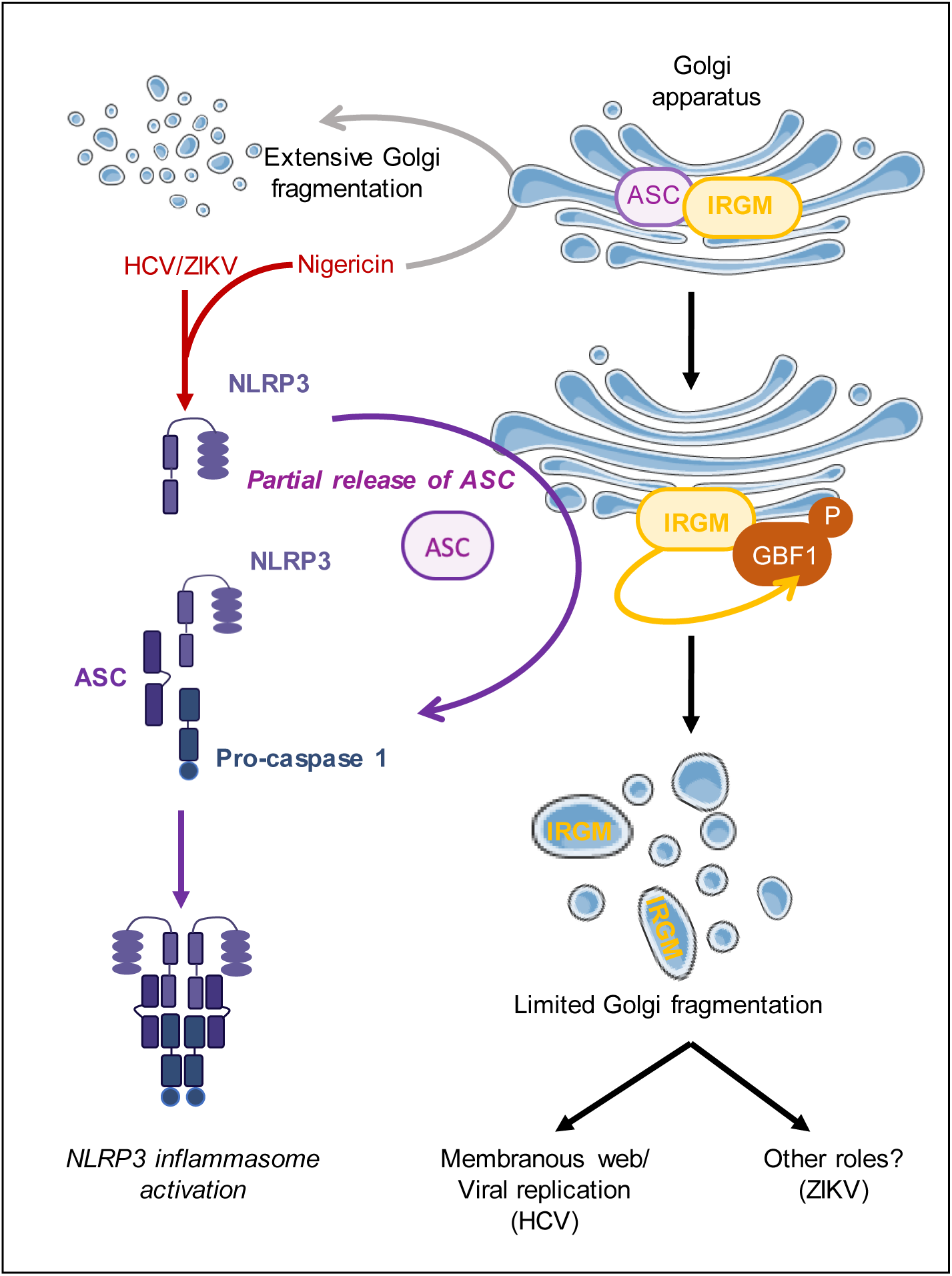
Proposed model for the role of NLRP3 and ASC in the IRGM-mediated Golgi fragmentation. Based on our results, we showed that ASC is a Golgi resident protein at homeostasis where it colocalizes with IRGM. HCV infection triggers induction of NLRP3 and we propose that this leads to a partial release of ASC from the Golgi, due to its association with NLRP3 in view to form an active NLRP3 inflammasome. Such release of ASC from the Golgi “liberates” IRGM which can then trigger Golgi fragmentation, through GBF1 phosphorylation, as reported previously (8). A link between Golgi fragmentation and NLRP3 can also occur after Zika virus (ZIKV) infection or after treatment with the potassium ionophore Nigericin. An NLRP3-independent extensive Golgi fragmentation was also noted in response to Nigericin. In contrast to HCV, ZIKV replication is reduced but not significantly decreased upon NLRP3 KO, despite the resulting reduced Golgi fragmentation. The role of Golgi fragmentation during ZIKV infection remains to be determined

The Golgi complex is organized in tubular structures in association with microtubules at a perinuclear localization. Although it is currently represented as a crescent-like twisted ribbon structure, the Golgi apparatus is now recognized as a very dynamic structure, which can adopt different configurations depending on the state of the cells (Makhoul, Gosavi, & Gleeson, 2018). This structure is maintained through the action of several Golgi resident proteins such as GBF1 (Saenz et al., 2009) and the GRASP proteins (Golgi Reassembly And Stacking Proteins) (Cervigni et al., 2015; Cheng et al., 2010). Alteration of this structure can occur in different situations, resulting notably in Golgi fragmentation. For instance, during the G2 phase of mitosis, JNK2 phosphorylates GRASP65, a protein that links the different Golgi cisternae, conveniently leading to equal transfer of the isolated Golgi stacks to the daughter cells (Cervigni et al., 2015). GBF1, a member of the Guanine Exchange Factor (GEF) family, maintains the integrity of the Golgi and coordinates the bidirectional transport of the COP proteins through the small GTPase Arf1. Phosphorylation of GBF1 allows the binding of Arf1 under its GTP-binding form to the Golgi, resulting in Golgi fragmentation. The modulation of GBF1 activity is observed during several infections, such as the enteroviruses poliovirus and coxsackievirus B3, for which GBF1 is crucial for viral replication (van der Linden, van der Schaar, Lanke, Neyts, & van Kuppeveld, 2010). Recently, we showed that HCV triggers GBF1 phosphorylation, and subsequent Golgi fragmentation through IRGM (Hansen et al., 2017). This process also benefits to HCV replication, as Golgi vesicular membranes can migrate to the viral replication complex where they can be used as lipid supply.

There are increasing numbers of reports showing inflammasome activation upon viral infections directly within the infected cells as well as in the surrounding immune cells. For instance, respiratory syncytial virus (RSV) activates the NLRP3 inflammasome by mediating the transfer of NLRP3 to the Golgi via its viral SH viroporin (Triantafilou, Kar, Vakakis, Kotecha, & Triantafilou, 2013). Recent reports on ZIKV showed that NS5 and NS1 viral proteins bind NLRP3 (Wang et al., 2018) and caspase-1 (Zheng et al., 2018), respectively. Inflammasome activation is well known to lead to inflammatory cytokine production (e.g., IL1ß, IL18) by immune cells and inflammation process. ZIKV infection leads to production of IL1ß by infected cells (Zheng et al., 2018), in contrast, activation of the inflammasome in HCV-infected cells did not cause IL1ß release, yet activation of inflammasome in surrounding cells could still be observed, presumably through the release of some DAMPs (Kofahi et al., 2016), in line with the observed severe liver inflammation and fibrogenesis, known to be associated with chronic Hepatitis C.

Very recent work suggested key functions for the Golgi apparatus and the ER-Golgi vesicle trafficking in the assembly of the NLRP3 inflammasome (Chen & Chen, 2018; C. Guo et al., 2018; Hong et al., 2019) and that disruption of Golgi integrity may inhibit NLRP3 inflammasome activation or, on the contrary, serves as a scaffold for NLRP3 aggregation and activation (Chen & Chen, 2018). This suggests that mediators from these organelles may contribute to inflammasome assembly.

Our data highlight that inflammasome components are utilized by the viruses in order to generate Golgi-derived lipids as supply for viral replication and assembly. HCV triggers the formation of a so-called membranous web close to the ER (Nagy & Pogany, 2012; Romero-Brey et al., 2012) while ZIKV triggers the remodeling of membranes, with apparition of convoluted membrane regions at the level of the smooth ER, for its replication (Rossignol et al., 2017). The formation of these structures requires the presence of lipids, which can be generated on site or delivered through transportation, such as provided through Golgi fragmentation. Thus, induction of the NLRP3 by HCV benefits the virus as it can lead to destabilization of the Golgi structure, thus allowing IRGM to further ensure Golgi fragmentation, allowing a supply of Golgi-derived lipids for HCV replication. It is possible to hypothesize that such prolonged expression of NLRP3 in hepatocytes, might be deleterious in the long term for the liver.

In the case of ZIKV, we showed that this virus can trigger Golgi fragmentation through NLRP3 such as HCV. However, in contrast to HCV, ZIKV replication is reduced but not significantly decreased upon NLRP3 KO, despite the resulting reduced Golgi fragmentation. We propose that Golgi-derived supply of lipids is not a limiting feature for ZIKV replication as compared to HCV and that ZIKV may use alternative mechanisms such as autophagy to provide lipids for its replication, as shown for other flaviviruses (Heaton & Randall, 2010).

While this work was in progress, IRGM has been reported to interact with ASC (Mehto et al., 2019). Interestingly, in this study, the IRGM/ASC interaction was observed in the inflammasome specks in hematopoietic cells submitted to different microbial PAMPs, and the data showed that IRGM was interrupting the interaction between NLRP3 and ASC, thus negatively controlling the NLRP3 inflammasome. On the other hand, we showed that IRGM and ASC interact at the Golgi in homeostatic conditions, and that they separate upon the presence and presumably activation of NLRP3. Thus, it is possible that the interaction of ASC and IRGM is essential to control the activity of each of these proteins. ASC retains IRGM at the Golgi and prevent its ability to trigger Golgi fragmentation as well as its ability to trigger autophagy while the association of IRGM to the inflammasome specks exerts a feedback control on the inflammasome by interrupting the NLRP3/ASC interaction. In line with this, the depletion of CPTP (Ceramide-1-phosphate transfer protein), involved in the traffic of Ceramide 1 phosphate from the trans-Golgi network to membranes, triggers Golgi fragmentation, activation of autophagy and of the NLRP3 inflammasome (Mishra et al., 2018). Future investigation will be needed to determine the effect of silencing of IRGM on Golgi structure, autophagy and inflammasome in this context.

The immunity-related GTPase M (IRGM) was previously identified as a risk factor for Crohn’s disease and tuberculosis, and for its role in autophagy. Data from Mehto et al. (Mehto et al., 2019) provided a mechanistic explanation for the ability of IRGM to control the inflammasome process through its interaction with ASC at the NLRP3 inflammasome and dragging the inflammasome to autophagosomes. Reciprocally, our data now provided a mechanistic explanation for the ability of the NLRP3 and ASC components of the inflammasome to trigger the IRGM-mediated Golgi fragmentation and autophagy. Golgi structure is known to be altered in stress related to disease (Bexiga & Simpson, 2013). In particular, a dispersed Golgi phenotype has been observed in several cancer cell lines and may play a central role in metastasis (Baschieri et al., 2014; Baschieri, Uetz-von Allmen, Legler, & Farhan, 2015; Kellokumpu, Sormunen, & Kellokumpu, 2002; Petrosyan, Holzapfel, Muirhead, & Cheng, 2014). Additionally, in Alzheimer’s disease, the Golgi is known to undergo fragmentation, and it is thought that fortifying Golgi stability could be a potential therapeutic strategy (Jiang et al., 2015; Joshi, Chi, Huang, & Wang, 2014; Stieber, Mourelatos, & Gonatas, 1996). Here, we uncovered that Golgi fragmentation, in conditions other than mitosis and cell division, can be considered as an important regulatory aspect of several pathological situations arising from viral infections. It should be therefore of high interest to examine the state of IRGM polymorphism in some chronic infections.

## Materials and Methods

### Cell Culture and virus

The hepatoma cell lines Huh7 (Arnaud et al., 2010) were maintained in DMEM supplemented with 10 % Fetal Bovine Serum (FBS) (Sigma, F7424), 1 % Penicillin/Streptomycin (Life Technologies) and 1 % non-essential amino acids (Sigma) (thereafter referred as complete medium) and cultured at 37 °C in 5 % CO2 The Huh7.25/CD81 and the Huh7.25/CD81/sgIRGM or Huh7.25/CD81/sgRenilla clones (Hansen et al., 2017) were maintained in the same medium with addition of 0.4 mg/ml geneticin (G418) for all cells and 10 µg/ml puromycin for the two latter ones. Macrophages were derived from monocytic U937 cells, which were KO or not (parental; WT) for NLRP3. U937 cells (obtained from the biological resource center, Cellulonet) were genetically modified by CRISPR/cas9 approach for the invalidation of NLRP3, as previously described (Lagrange et al., 2018). U937 cells and derivates were cultured in RPMI 1640 supplemented with 10 % FBS, 100 IU/mL penicillin, 100 μg/mL streptomycin (all from Life Technologies) at 37 °C in 5 % CO2 and differentiated into macrophage-like phenotype cells by adding 100 nM of phorbol 12-myristate 13-acetate (PMA; InvivoGen) two days prior infection by ZIKV.

JFH1 HCV stocks were produced in Huh7.25/CD81 cells and titrated as described (Arnaud et al., 2010). HCV infections were performed with a multiplicity of infection (MOI) of 0.3 in complete medium for 2 h before the removal of virus and incubation for the indicated time points. For 5 days infection, cells were trypsinized and split at day 3 PI before further cell expansion for the 2 remaining days. The clinical isolate of Brazil strain of ZIKV: PE243_KX197192) (Donald et al., 2016) was amplified in Vero cells. The supernatant was collected three days post-infection, filtered with 0.45 μ-m filter (Corning) and stored at −80°C. Prior to infection of macrophages from monocytic U937 cells KO for NLRP3 and parental cells, ZIKV was incubated with 4G2 antibody (1:100 dilution; hybridoma supernatant-containing anti-E glycoprotein of flavivirus) for one hour at 37°C. ZIKV infections were performed at MOI of 5 for U937 cells for the indicated time points.

### Reagents and Antibodies

Primary antibodies were purchased from Santa Cruz: HCV core (mouse mAb, sc-57800), ASC (mouse mAb, sc-514414); from Cell Signaling Technologies: NLRP3 (rabbit mAb, 15101S), normal rabbit IgG (2729); from Abcam: IRGM (mouse mAb, ab-69494), GM130 (rabbit mAb 52649); from Sigma: GAPDH (mouse mAb, g8795) and from BD transduction: GM130 (mouse Mab, 610822), total GBF1 (mouse mAb, 612116); from IBL: P-GBF1 (T1337 Phosphorylated) (rabbit Ab, 28065); from Adipogen: NLRP3 (mouse mAb, AG-20B-0014), ASC (rabbit, AG-25B-0006). The hybridoma producing antibodies against Flavivirus Envelope protein 4G2 was kindly provided by P. Despres (PIMIT, Université de La Réunion-INSERM France). Secondary antibodies for immunoblotting were from Thermo Scientific (Goat anti-Mouse IgG DyLight 800 and DyLight 680, Goat anti Rabbit IgG DyLight 800 and DyLight 680). Secondary antibodies for immunofluorescence were from Invitrogen (Alexa Fluor 488 Goat anti-Mouse and Goat anti-Rabbit, Alexa Fluor 568 Goat anti-Mouse and Goat anti-Rabbit). Anti-proteases cOmplete and anti-phosphatases PhoSTOP were from Roche. Nigericin was from Sigma. Caspase-1 activation was detected using the FAM-FLICA® Caspase1 assay kit (ImmunoChemistry Technologies). Previous reagents and antibodies were used following the manufacturer’s instructions.

### RNAi. The following siRNA oligonucleotides were used

NLRP3 (Dharmacon; 114548), ASC (Qiagen; 1027416), Ambion AM51331, siRNA ID: 44415 (siASC 2) Ambion 4392420, siRNA ID: s26508 (siASC 3), negative control siRNA (Dharmacon; D-001810-10-20). siRNA duplexes were reverse transfected into cells using Lipofectamine RNAiMAX (Invitrogen;17-0618-0) at the day of plating (24hr before infection with HCV) according to the manufacturer’s instructions. Two subsequent transfections were performed on adherent cells, the first one a few hours before infection and the second one, at day 3 post infection.

### Quantitative real-time PCR analysis

Total cellular RNA was extracted using TRI-Reagent (Sigma), according to the manufacturer’s instructions. cDNA was generated from 1 µg of total RNA using the Revert Aid reagents (ThermoScientific). The resulting cDNA was diluted 1:20 and subjected to semi-quantitative real time PCR using the SYBR Green reagent (Life Technologies) and specific primers. Those primers were: JFH1, 5’-TCCCGGGAGAGCCATAGTG-3’ (forward) and 5’-TCCAGGAAAGGACCCAGTC-3’ (reverse). Roth, 5’-CTGCGGACTATCTCTCCCCTC-3’ (forward) and 5’-AAAAGGCTTTGCAGCTCCAC-3’ (reverse). ZIKV, 5’-ATTGTTGGTGCAACACGACG-3’ (forward) and 5’-CCTAGTGGAATGGGAGGGGA-3’ (reverse). IL8, 5’-AAGGGCCAAGAGAATATCCGAA-3’ (forward) and 5’-ACTAGGGTTGCCAGATTTAACA-3’ (reverse). Amplification reactions were performed under the following conditions: 2 min at 50 °C, 10 min at 95 °C, 40 cycles for 10 s at 95 °C, and 1 min at 60 °C. As specified by the manufacturer, relative transcript levels were calculated using the ΔCt method, using Roth as the reference gene.

### Immunofluorescence

Cells were fixed in PBS containing 4% paraformaldehyde (PFA) for 15 min on ice and were permeabilized and non-specific antibody sites were blocked with PBS containing 10 % FBS, 2,5 % BSA and 0,3 % Triton-X100 for 30 min at room temperature. For staining, cells were incubated with antibodies diluted in blocking solution. For U937 cells, human Fc blocking reagent (MACS Miltenyi Biotec) was added to this blocking step. Primary antibodies were added for 60 min at 37°C or at 4°C overnight. Secondary antibodies were added for 30 min at room temperature. Nuclei were visualized by incubation with DAPI for 5 min at room temperature. For staining IRGM proteins, cells were fixed and permeabilized with 100% ice cold Methanol at −20°C for 10 min and blocked with PBS containing 10 % FBS and 2,5 % BSA for 30 min at room temperature.

### Confocal Fluorescence Microscopy

Confocal fluorescence microscopy studies were performed with a Zeiss Axiovert 200-M inverted microscope equipped with an LSM 510 or 700 laser-scanning unit and a 1.4 NA, 63x Plan-Apochromat oil-immersion objective. Cells were seeded in µ-Slide eight-well, ibiTreat tissue culture-treated plates (Inter Instrument AS), in 12-well plates containing glass coverslips (Sigma) or in 96-Well glass bottom plates (Nunc). The glass coverslips were mounted with Prolong (Invitrogen). To minimize photobleaching, laser power typically was 20 % under maximum, and the pinhole was set to 0.8 – 1.5. Multitracking was used for dual- or triple-color imaging. Quantitative confocal image analysis was performed using ImageJ software. The fluorescence intensity of Golgi objects (based on GM130 staining) was determined by background subtraction and using a fixed threshold. Otsu’s thresholding algorithm was applied to convert images to binary images and to create ROIs based on pixel intensity. Then the particle area and circularity was quantified using the Analyze Particle plugin in ImageJ (https://imagej.nih.gov/ij)). A circularity value of 1 corresponds to a perfect circle (Miller et al., 2009). Colocalization analysis was performed using the JaCoP plugin in ImageJ (Bolte & Cordelieres, 2006) Image parameters remained constant during imaging, and threshold values were kept the same from image to image during image analysis. Mander’s colocalization coefficient (MCC) was used. MCCs measure the fraction of one protein that colocalizes with a second protein (Dunn, Kamocka, & McDonald, 2011).

### Proximity Ligation Assay (PLA)

Cells were cultured on 24-well plates containing glass coverslips, washed with PBS, fixed and permeabilized with Aceton/Methanol (isovolume) for 5 min at −20°C, except for the ASC/NLRP3 PLA, where the cells were fixed with paraformaldehyde 4 % (PFA) for 15 min at room temperature and permeabilized with digitonin 0,01% for 30 min at room temperature.

Proximity ligation assay was performed using the DuoLink PLA technology (Sigma). Briefly, the cells were incubated with Duolink blocking solution for 1 h at 37 °C in a pre-heated humidity chamber, and then with primary antibodies overnight at 4 °C. The coverslips were washed with buffer A and incubated with the PLUS and MINUS PLA probes (antibodies with attached oligonucleotide strands) for 1 h at 37 °C. After three washes with buffer A, the PLA probes were hybridized to connect oligonucleotides which are then joined in a ligation step for 30 min at 37 °C. After three more washes with buffer A, the resulting closed circular DNA template is amplified by DNA polymerase for 100 min at 37 °C. Oligos coupled to fluorochromes contained in the amplification buffer hybridize to repeated motifs on the amplicons, allowing detection. Coverslips were then washed with buffer B and mounted with Duolink *in situ* mounting medium containing DAPI. Images were collected (Zeiss Inverted LSM 700 Microscope) and analyzed with ImageJ: a projection of Z-stack PLA blobs was merged with nuclei labeled with DAPI.

### Immunoblotting

Cells were washed once with PBS and lysed either in CHAPS lysis buffer containing 50 mM Tris (pH 7.5), 140 mM NaCl, 5% glycerol, 1% CHAPS, 2 mM EDTA, 40 mM glycerophosphate, or in 8M Urea buffer, both containing protease and phosphatase inhibitors (cOmplete inhibitor cocktail; Roche). Cell lysates were clarified by centrifugation, separated by lithium dodecyl sulfate (LDS)-PAGE, and electrophoretically transferred to nitrocellulose membranes using iBlot transfer system (Invitrogen, Life Technologies). Membranes were blocked in Li-Cor Odyssey blocking buffer at 1:1 ratio with Tris-buffered saline (TBS) for 1 hr at room temperature before incubation with the indicated antibodies overnight a 4°C. After washes with TBS containing 0,1 % Tween 20, immunoreactive proteins were detected with Li-COR DyLight secondary antibodies and visualized on the Li-COR Odyssey system.

### Statistical analysis

For quantification of immunofluorescence microscopy images, a minimum of 20 cells was counted for each condition in each experiment. Three independent experiments were performed for all figures. Data are shown as mean ± SD. P values were calculated by using paired Student t test, and a P value <0.05 was considered statistically significant. *, P<0.05; **, P<0.01; ***, P<0.001.

## Acknowledgements

We thank Alain Kohn (MRC-University of Glasgow Centre for Virus Research, UK) and Rafael Freitas de Oliveira Franca (FIOCRUZ, Recife, Brazil) for clinical isolate Brazil, (PE243_KX197192) and P. Despres (PIMIT, Université de La Réunion-INSERM France)) for providing the pan-flavirus antibody (4G2). Confocal imaging was performed at the Cellular and Molecular Imaging Core Facility at NTNU and at the Photonic BioImaging Unit of Technology and Service at Institut Pasteur. This work was supported by grants from The Liaison Committee for Education, Research and Innovation in Central Norway, to MDH, the ‘Agence Nationale pour la Recherche contre le SIDA et les Hépatites Virales’ to EM and MD, the ‘Agence Nationale pour la Recherche’ (ANR-JCJC) and LabEx Ecofect Grant ANR-11-LABX-0048 to MD.

## References

Amako, Y., Syed, G. H., & Siddiqui, A. (2011). Protein kinase D negatively regulates hepatitis C virus secretion through phosphorylation of oxysterol-binding protein and ceramide transfer protein. J Biol Chem, 286 (13), 11265–11274. doi:10.1074/jbc.M110.182097

Arnaud, N., Dabo, S., Maillard, P., Budkowska, A., Kalliampakou, K. I., Mavromara, P., … Meurs, E. F. (2010). Hepatitis C virus controls interferon production through PKR activation. PLoS One, 5 (5), e10575. doi:10.1371/journal.pone.0010575 [doi]

Baschieri, F., Confalonieri, S., Bertalot, G., Di Fiore, P. P., Dietmaier, W., Leist, M., … Farhan, H. (2014). Spatial control of Cdc42 signalling by a GM130-RasGRF complex regulates polarity and tumorigenesis. Nat Commun, 5, 4839. doi:10.1038/ncomms5839

Baschieri, F., Uetz-von Allmen, E., Legler, D. F., & Farhan, H. (2015). Loss of GM130 in breast cancer cells and its effects on cell migration, invasion and polarity. Cell Cycle, 14 (8), 1139–1147. doi:10.1080/15384101.2015.1007771

Bexiga, M. G., & Simpson, J. C. (2013). Human diseases associated with form and function of the Golgi complex. Int J Mol Sci, 14 (9), 18670–18681. doi:10.3390/ijms140918670

Bolte, S., & Cordelieres, F. P. (2006). A guided tour into subcellular colocalization analysis in light microscopy. J Microsc, 224 (Pt 3), 213–232. doi:10.1111/j.1365-2818.2006.01706.x

Cervigni, R. I., Bonavita, R., Barretta, M. L., Spano, D., Ayala, I., Nakamura, N., … Colanzi, A. (2015). JNK2 controls fragmentation of the Golgi complex and the G2/M transition through phosphorylation of GRASP65. J Cell Sci, 128 (12), 2249–2260. doi:10.1242/jcs.164871

Chauhan, S., Mandell, M. A., & Deretic, V. (2016). Mechanism of action of the tuberculosis and Crohn disease risk factor IRGM in autophagy. Autophagy, 12 (2), 429–431. doi:10.1080/15548627.2015.1084457

Chen, J., & Chen, Z. J. (2018). PtdIns4P on dispersed trans-Golgi network mediates NLRP3 inflammasome activation. Nature, 564 (7734), 71–76. doi:10.1038/s41586-018-0761-3

Cheng, J. P., Betin, V. M., Weir, H., Shelmani, G. M., Moss, D. K., & Lane, J. D. (2010). Caspase cleavage of the Golgi stacking factor GRASP65 is required for Fas/CD95-mediated apoptosis. Cell Death Dis, 1, e82. doi:10.1038/cddis.2010.59

Chusri, P., Kumthip, K., Hong, J., Zhu, C., Duan, X., Jilg, N., … Chung, R. T. (2016). HCV induces transforming growth factor beta1 through activation of endoplasmic reticulum stress and the unfolded protein response. Sci Rep, 6, 22487. doi:10.1038/srep22487

Donald, C. L., Brennan, B., Cumberworth, S. L., Rezelj, V. V., Clark, J. J., Cordeiro, M. T., … Kohl, A. (2016). Full Genome Sequence and sfRNA Interferon Antagonist Activity of Zika Virus from Recife, Brazil. PLoS Negl Trop Dis, 10 (10), e0005048. doi:10.1371/journal.pntd.0005048

Dunn, K. W., Kamocka, M. M., & McDonald, J. H. (2011). A practical guide to evaluating colocalization in biological microscopy. Am J Physiol Cell Physiol, 300 (4), C723–742. doi:10.1152/ajpcell.00462.2010

Goossens, N., & Hoshida, Y. (2015). Hepatitis C virus-induced hepatocellular carcinoma. Clin Mol Hepatol, 21 (2), 105–114. doi:10.3350/cmh.2015.21.2.105

Gregoire, I. P., Richetta, C., Meyniel-Schicklin, L., Borel, S., Pradezynski, F., Diaz, O., … Faure, M. (2011). IRGM is a common target of RNA viruses that subvert the autophagy network. PLoS Pathog, 7 (12), e1002422. doi:10.1371/journal.ppat.1002422

Guo, C., Chi, Z., Jiang, D., Xu, T., Yu, W., Wang, Z., … Wang, D. (2018). Cholesterol Homeostatic Regulator SCAP-SREBP2 Integrates NLRP3 Inflammasome Activation and Cholesterol Biosynthetic Signaling in Macrophages. Immunity, 49 (5), 842-856.e847. doi:10.1016/j.immuni.2018.08.021

Guo, H., Callaway, J. B., & Ting, J. P. (2015). Inflammasomes: mechanism of action, role in disease, and therapeutics. Nat Med, 21 (7), 677–687. doi:10.1038/nm.3893

Hansen, M. D., Johnsen, I. B., Stiberg, K. A., Sherstova, T., Wakita, T., Richard, G. M., … Anthonsen, M. W. (2017). Hepatitis C virus triggers Golgi fragmentation and autophagy through the immunity-related GTPase M. Proc Natl Acad Sci U S A, 114 (17), E3462–e3471. doi:10.1073/pnas.1616683114

Heaton, N. S., & Randall, G. (2010). Dengue virus-induced autophagy regulates lipid metabolism. Cell Host Microbe, 8 (5), 422–432. doi:10.1016/j.chom.2010.10.006

Hong, S., Hwang, I., Gim, E., Yang, J., Park, S., Yoon, S. H., … Yu, J. W. (2019). Brefeldin A-sensitive ER-Golgi vesicle trafficking contributes to NLRP3-dependent caspase-1 activation. Faseb j, 33 (3), 4547–4558. doi:10.1096/fj.201801585R

Jiang, Y., Bao, H., Ge, Y., Tang, W., Cheng, D., Luo, K., … Gong, R. (2015). Therapeutic targeting of GSK3beta enhances the Nrf2 antioxidant response and confers hepatic cytoprotection in hepatitis C. Gut, 64 (1), 168–179. doi:10.1136/gutjnl-2013-306043

Joshi, G., Chi, Y., Huang, Z., & Wang, Y. (2014). Abeta-induced Golgi fragmentation in Alzheimer’s disease enhances Abeta production. Proc Natl Acad Sci U S A, 111 (13), E1230–1239. doi:10.1073/pnas.1320192111

Kellokumpu, S., Sormunen, R., & Kellokumpu, I. (2002). Abnormal glycosylation and altered Golgi structure in colorectal cancer: dependence on intra-Golgi pH. FEBS Lett, 516 (1-3), 217–224. doi:10.1016/s0014-5793(02)02535-8

Kofahi, H. M., Taylor, N. G., Hirasawa, K., Grant, M. D., & Russell, R. S. (2016). Hepatitis C Virus Infection of Cultured Human Hepatoma Cells Causes Apoptosis and Pyroptosis in Both Infected and Bystander Cells. Sci Rep, 6, 37433. doi:10.1038/srep37433

Lagrange, B., Benaoudia, S., Wallet, P., Magnotti, F., Provost, A., Michal, F., … Henry, T. (2018). Human caspase-4 detects tetra-acylated LPS and cytosolic Francisella and functions differently from murine caspase-11. Nat Commun, 9 (1), 242. doi:10.1038/s41467-017-02682-y

Lavie, M., & Dubuisson, J. (2017). Interplay between hepatitis C virus and lipid metabolism during virus entry and assembly. Biochimie, 141, 62–69. doi:10.1016/j.biochi.2017.06.009

Lebsir, N., Goueslain, L., Farhat, R., Callens, N., Dubuisson, J., Jackson, C. L., & Rouille, Y. (2019). Functional and Physical Interaction between the Arf Activator GBF1 and Hepatitis C Virus NS3 Protein. J Virol, 93 (6). doi:10.1128/jvi.01459-18

Makhoul, C., Gosavi, P., & Gleeson, P. A. (2018). The Golgi architecture and cell sensing. Biochem Soc Trans, 46 (5), 1063–1072. doi:10.1042/bst20180323

Mariathasan, S., Weiss, D. S., Newton, K., McBride, J., O’Rourke, K., Roose-Girma, M., … Dixit, V. M. (2006). Cryopyrin activates the inflammasome in response to toxins and ATP. Nature, 440 (7081), 228–232. doi:10.1038/nature04515

Mehto, S., Jena, K. K., Nath, P., Chauhan, S., Kolapalli, S. P., Das, S. K., … Chauhan, S. (2019). The Crohn’s Disease Risk Factor IRGM Limits NLRP3 Inflammasome Activation by Impeding Its Assembly and by Mediating Its Selective Autophagy. Mol Cell, 73 (3), 429-445.e427. doi:10.1016/j.molcel.2018.11.018

Miller, P. M., Folkmann, A. W., Maia, A. R., Efimova, N., Efimov, A., & Kaverina, I. (2009). Golgi-derived CLASP-dependent microtubules control Golgi organization and polarized trafficking in motile cells. Nat Cell Biol, 11 (9), 1069–1080. doi:10.1038/ncb1920

Mirandola, S., Bowman, D., Hussain, M. M., & Alberti, A. (2010). Hepatic steatosis in hepatitis C is a storage disease due to HCV interaction with microsomal triglyceride transfer protein (MTP). Nutr Metab (Lond), 7, 13. doi:10.1186/1743-7075-7-13

Misawa, T., Takahama, M., Kozaki, T., Lee, H., Zou, J., Saitoh, T., & Akira, S. (2013). Microtubule-driven spatial arrangement of mitochondria promotes activation of the NLRP3 inflammasome. Nat Immunol, 14 (5), 454–460. doi:10.1038/ni.2550

Mishra, S. K., Gao, Y. G., Deng, Y., Chalfant, C. E., Hinchcliffe, E. H., & Brown, R. E. (2018). CPTP: A sphingolipid transfer protein that regulates autophagy and inflammasome activation. Autophagy, 14 (5), 862–879. doi:10.1080/15548627.2017.1393129

Nagy, P. D., & Pogany, J. (2012). The dependence of viral RNA replication on co-opted host factors. Nat Rev Microbiol, 10 (2), 137–149. doi:10.1038/nrmicro2692

Negash, A. A., & Gale, M., Jr. (2015). Hepatitis regulation by the inflammasome signaling pathway. Immunol Rev, 265 (1), 143–155. doi:10.1111/imr.12279

Petrosyan, A., Holzapfel, M. S., Muirhead, D. E., & Cheng, P. W. (2014). Restoration of compact Golgi morphology in advanced prostate cancer enhances susceptibility to galectin-1-induced apoptosis by modifying mucin O-glycan synthesis. Mol Cancer Res, 12 (12), 1704–1716. doi:10.1158/1541-7786.Mcr-14-0291-t

Reiss, S., Rebhan, I., Backes, P., Romero-Brey, I., Erfle, H., Matula, P., … Bartenschlager, R. (2011). Recruitment and activation of a lipid kinase by hepatitis C virus NS5A is essential for integrity of the membranous replication compartment. Cell Host Microbe, 9 (1), 32–45. doi:10.1016/j.chom.2010.12.002

Romero-Brey, I., & Bartenschlager, R. (2014). Membranous replication factories induced by plus-strand RNA viruses. Viruses, 6 (7), 2826–2857. doi:10.3390/v6072826

Romero-Brey, I., Merz, A., Chiramel, A., Lee, J. Y., Chlanda, P., Haselman, U., … Bartenschlager, R. (2012). Three-dimensional architecture and biogenesis of membrane structures associated with hepatitis C virus replication. PLoS Pathog, 8 (12), e1003056. doi:10.1371/journal.ppat.1003056

Rossignol, E. D., Peters, K. N., Connor, J. H., & Bullitt, E. (2017). Zika virus induced cellular remodelling. Cell Microbiol, 19 (8). doi:10.1111/cmi.12740

Saenz, J. B., Sun, W. J., Chang, J. W., Li, J., Bursulaya, B., Gray, N. S., & Haslam, D. B. (2009). Golgicide A reveals essential roles for GBF1 in Golgi assembly and function. Nat Chem Biol, 5 (3), 157–165. doi:10.1038/nchembio.144

Sagulenko, V., Thygesen, S. J., Sester, D. P., Idris, A., Cridland, J. A., Vajjhala, P. R., … Stacey, K. J. (2013). AIM2 and NLRP3 inflammasomes activate both apoptotic and pyroptotic death pathways via ASC. Cell Death Differ, 20 (9), 1149–1160. doi:10.1038/cdd.2013.37

Sanchez-San Martin, C., Li, T., Bouquet, J., Streithorst, J., Yu, G., Paranjpe, A., & Chiu, C. Y. (2018). Differentiation enhances Zika virus infection of neuronal brain cells. Sci Rep, 8 (1), 14543. doi:10.1038/s41598-018-32400-7

Song, N., Liu, Z. S., Xue, W., Bai, Z. F., Wang, Q. Y., Dai, J., … Li, T. (2017). NLRP3 Phosphorylation Is an Essential Priming Event for Inflammasome Activation. Mol Cell, 68 (1), 185-197.e186. doi:10.1016/j.molcel.2017.08.017

Spengler, U. (2018). Direct antiviral agents (DAAs) - A new age in the treatment of hepatitis C virus infection. Pharmacol Ther, 183, 118–126. doi:10.1016/j.pharmthera.2017.10.009

Stieber, A., Mourelatos, Z., & Gonatas, N. K. (1996). In Alzheimer’s disease the Golgi apparatus of a population of neurons without neurofibrillary tangles is fragmented and atrophic. Am J Pathol, 148 (2), 415–426.

Sun, B., & Karin, M. (2008). NF-kappaB signaling, liver disease and hepatoprotective agents. Oncogene, 27 (48), 6228–6244. doi:10.1038/onc.2008.300

Triantafilou, K., Kar, S., Vakakis, E., Kotecha, S., & Triantafilou, M. (2013). Human respiratory syncytial virus viroporin SH: a viral recognition pathway used by the host to signal inflammasome activation. Thorax, 68 (1), 66–75. doi:10.1136/thoraxjnl-2012-202182

van der Linden, L., van der Schaar, H. M., Lanke, K. H., Neyts, J., & van Kuppeveld, F. J. (2010). Differential effects of the putative GBF1 inhibitors Golgicide A and AG1478 on enterovirus replication. J Virol, 84 (15), 7535–7542. doi:10.1128/jvi.02684-09

Wang, W., Li, G., De, W., Luo, Z., Pan, P., Tian, M., … Wu, J. (2018). Zika virus infection induces host inflammatory responses by facilitating NLRP3 inflammasome assembly and interleukin-1beta secretion. Nat Commun, 9 (1), 106. doi:10.1038/s41467-017-02645-3

Zhang, Z., Meszaros, G., He, W. T., Xu, Y., de Fatima Magliarelli, H., Mailly, L., … Ricci, R. (2017). Protein kinase D at the Golgi controls NLRP3 inflammasome activation. J Exp Med, 214 (9), 2671–2693. doi:10.1084/jem.20162040

Zheng, Y., Liu, Q., Wu, Y., Ma, L., Zhang, Z., Liu, T., … Cui, J. (2018). Zika virus elicits inflammation to evade antiviral response by cleaving cGAS via NS1-caspase-1 axis. Embo j, 37 (18). doi:10.15252/embj.201899347

Zhou, R., Yazdi, A. S., Menu, P., & Tschopp, J. (2011). A role for mitochondria in NLRP3 inflammasome activation. Nature, 469 (7329), 221–225. doi:nature09663 [pii] 10.1038/nature09663 [doi]

