## Supplementary figures and legends for "The inflammasome components NLRP3 and ASC act in concert with IRGM to rearrange the Golgi during viral infections"

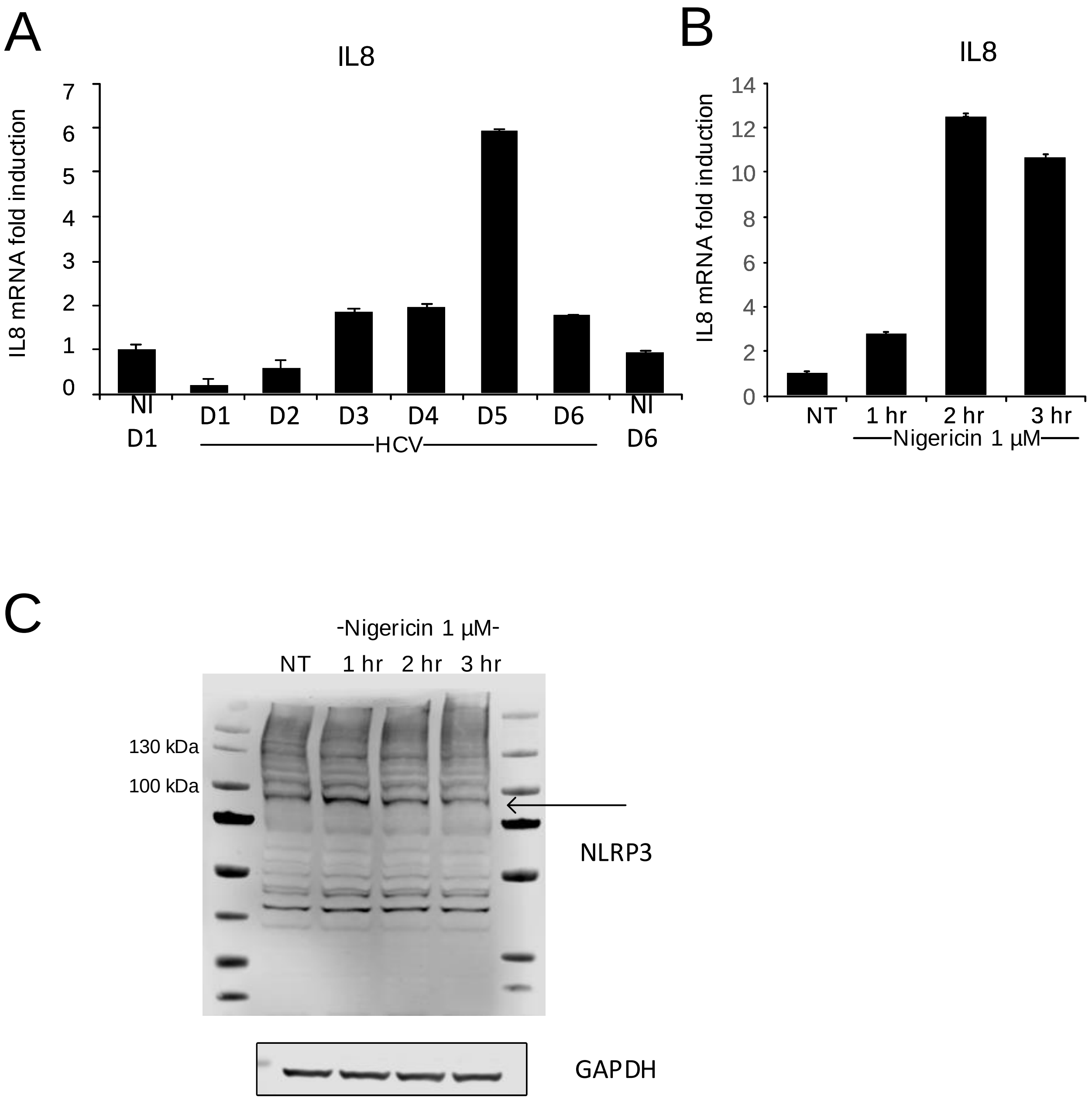
A
IL8
IL8 mRNA fold induction
7
6
5
4
3
2
1
0
NI D1
D1
D2
D3
D4
D5
D6
NI D6
HCV
B
IL8
IL8 mRNA fold induction
14
12
10
8
6
4
2
0
NT
1 hr
2 hr
3 hr
Nigericin 1 µM
C
-Nigericin 1 µM-
NT
1 hr
2 hr
3 hr
130 kDa
100 kDa
NLRP3
GAPDH

**SFigure 1**

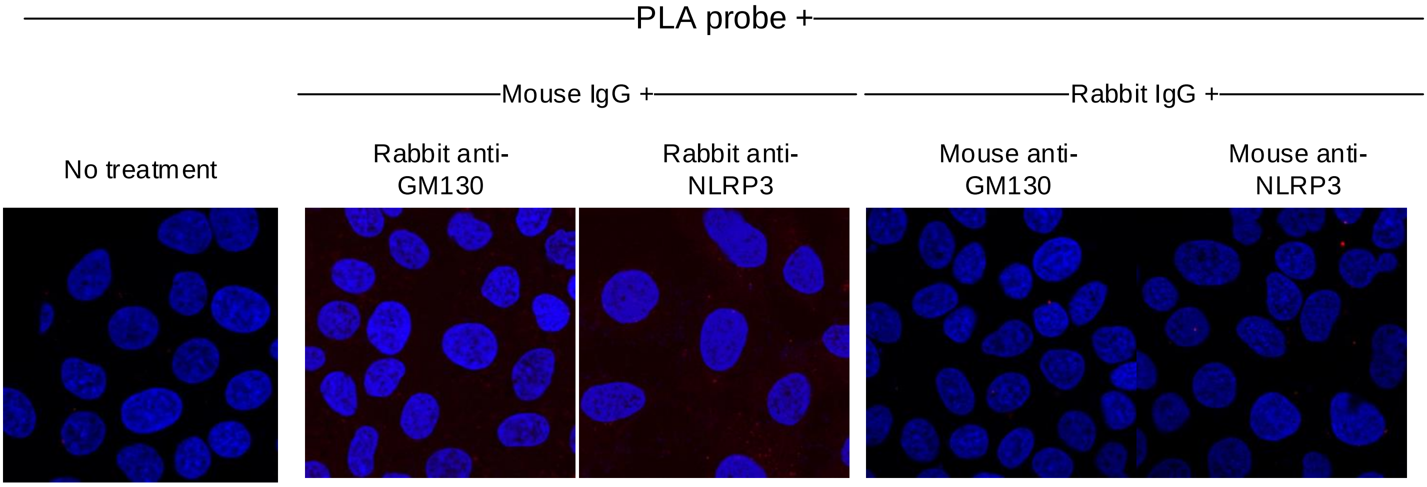
PLA probe +
Mouse IgG +
Rabbit IgG +
No treatment
Rabbit anti-GM130
Rabbit anti-NLRP3
Mouse anti-GM130
Mouse anti-NLRP3

**SFigure 2**

A

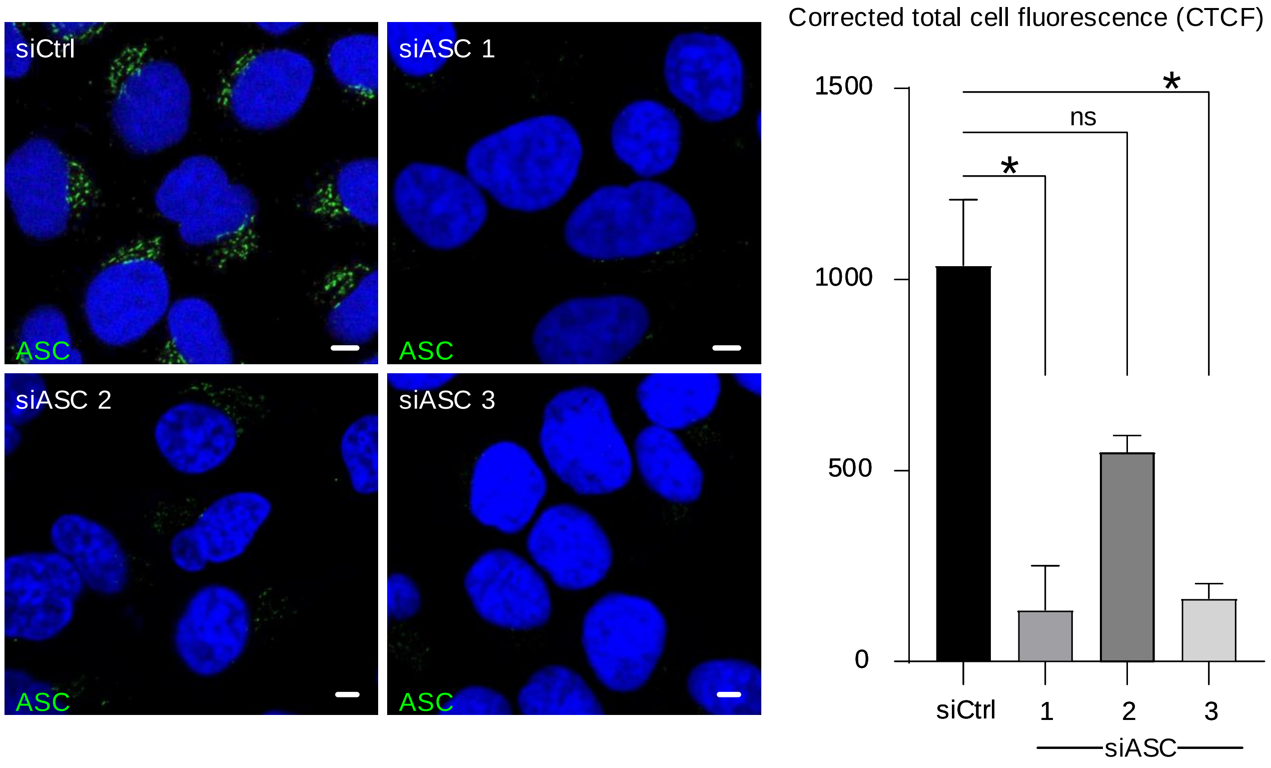

siCtrl
siASC 1
siASC 2
siASC 3
ASC
Corrected total cell fluorescence (CTCF)
1500
1000
500
0
ns
*
siCtrl
1
2
3
siASC


B

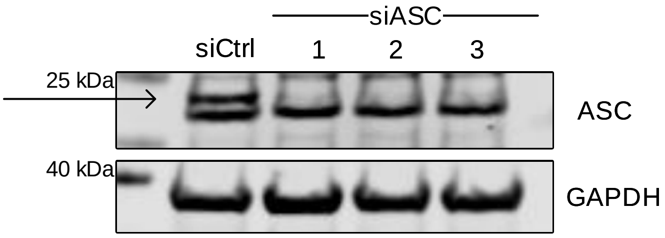

siASC
siCtrl
1
2
3
25 kDa
40 kDa
ASC
GAPDH


**Sfigure 3**

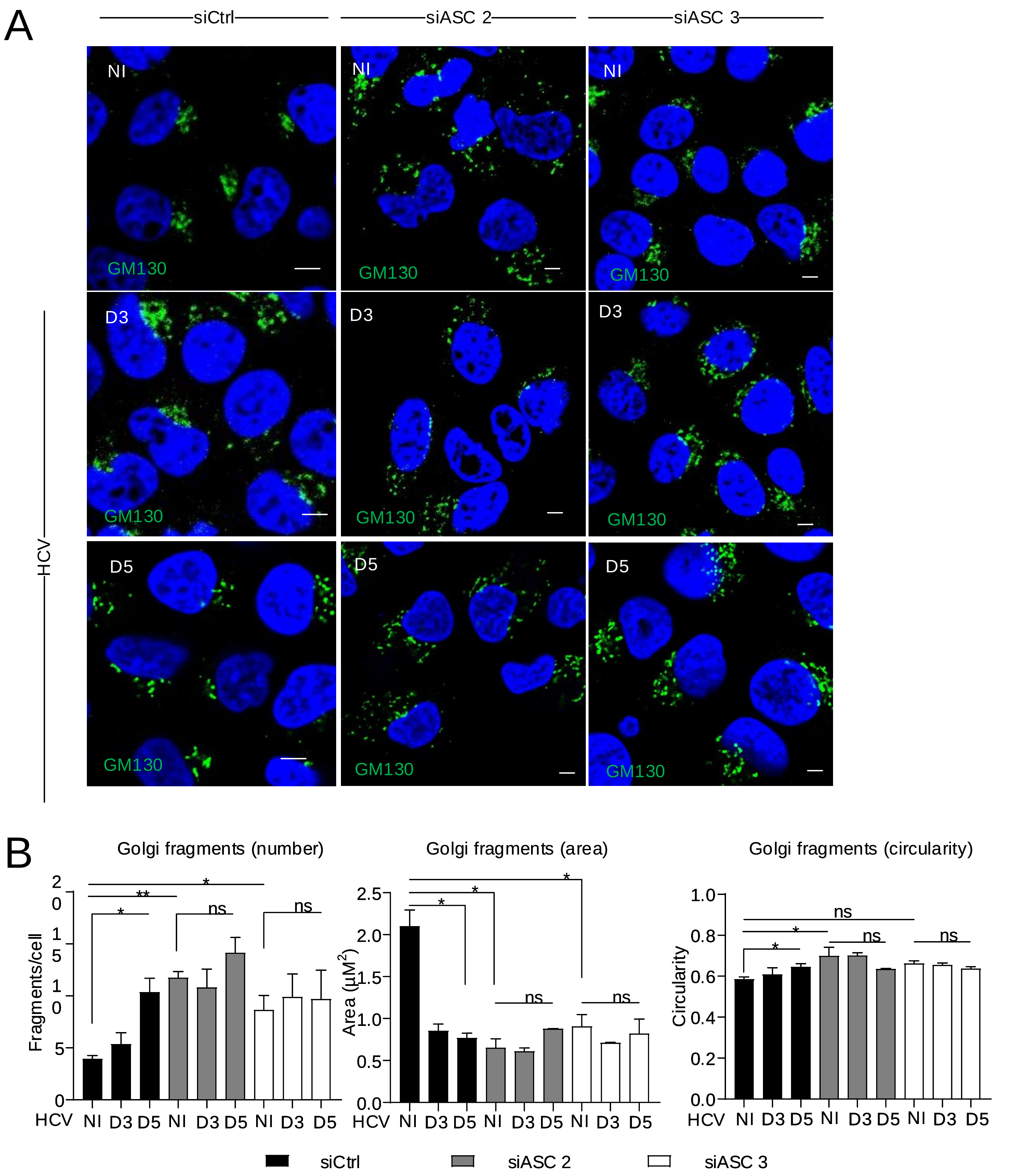

A
siCtrl
siASC 2
siASC 3
NI
D3
D5
GM130
HCV
B
Golgi fragments (number)
Golgi fragments (area)
Golgi fragments (circularity)
Fragments/cell
Area (μM2)
Circularity
HCV NI D3 D5 NI D3 D5 NI D3 D5
ns
siCtrl
siASC 2
siASC 3


**Sfigure 4**

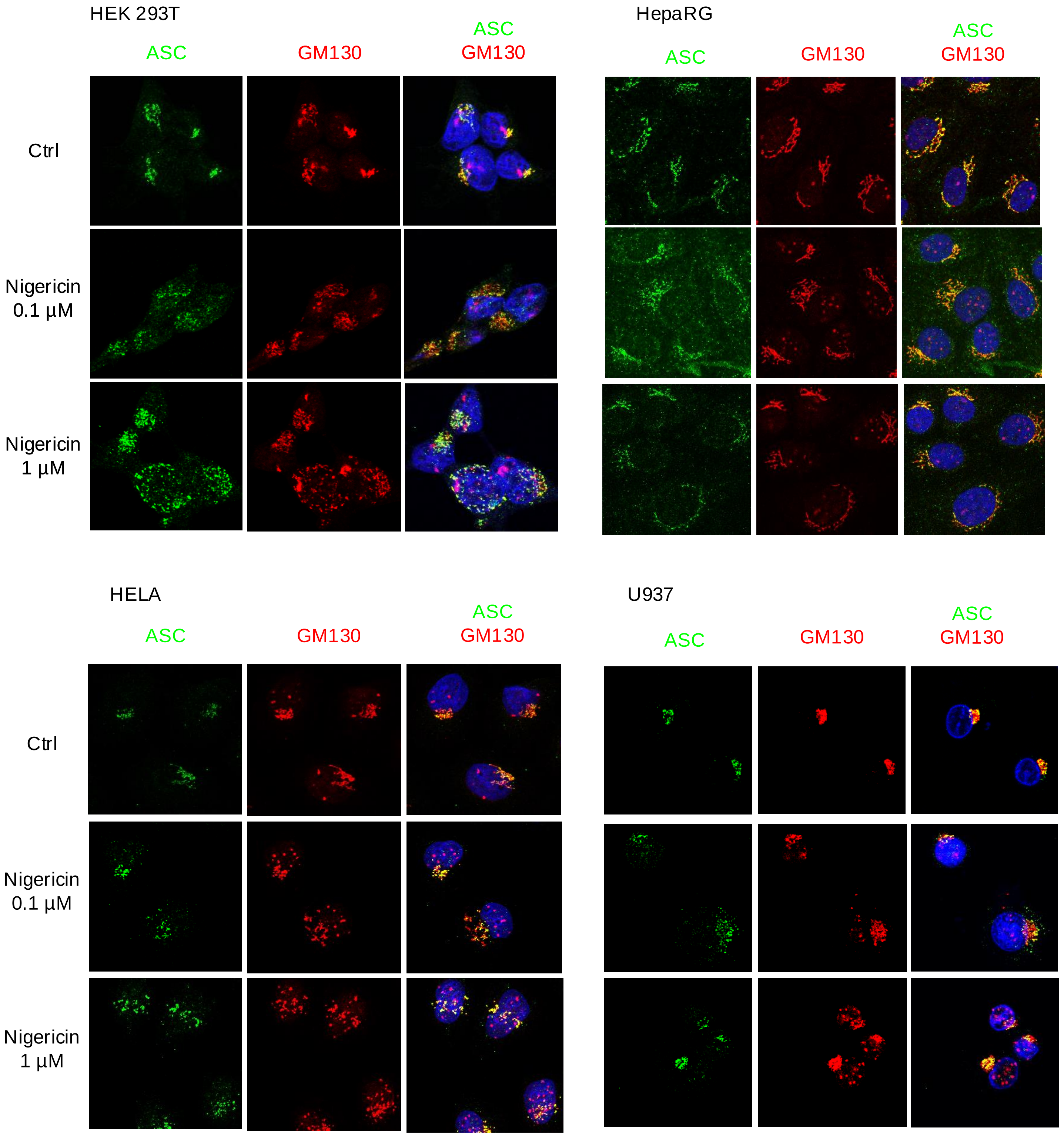
HEK 293T
ASC
GM130
ASC
GM130
Ctrl
Nigericin
0.1 µM
Nigericin
1 µM
HepaRG
ASC
GM130
ASC
GM130
HELA
ASC
GM130
ASC
GM130
Ctrl
Nigericin
0.1 µM
Nigericin
1 µM
U937
ASC
GM130
ASC
GM130

**Sfigure 5**

### Supplementary figure legends

**SFig 1:**

(A) IL8 mRNA levels were determined by qRTPCR in cells non-infected or infected with HCV for 1 to 6 days PI. Fold change values have been calculated relative to non-infected day 1 (B) IL8 mRNA levels were determined by qRTPCR in cells treated with Nigericin (1 µM; 1, 2 and 3 hours of treatment). Fold change values have been calculated relative to non-treated condition. (C) The effect of Nigericin on NLRP3 protein levels was examined by immunoblotting. A blot from one representative experiment is shown. Cells were treated with Nigericin (1 µM; 1, 2 and 3 hours of treatment) or left untreated. GAPDH was used as a loading control. Arrow is marking the NLRP3 band.

**SFig 2: PLA controls**

Representative images of PLA performed on controls. Cells were left untreated or incubated with each of the antibodies used in this assay. Verifying the specificity of the assay.

**SFig 3: Control of ASC knockdown**

Huh7 cells were treated with three different siRNAs against ASC or Control siRNA. (A) Intracellular staining of endogenous ASC. Cells were examined by confocal microscopy and the corrected total cell fluorescence (CTCF) were calculated using ImageJ software (B) Total protein levels were examined by immunoblotting with an ASC antibody. The migration of the molecular weight markers is presented to point out that the anti-ASC antibody recognize specifically ASC at its expected position. The specificity of the siASCs was verified, as they silenced only the target gene and did not affect intensity of non-specific bands.

**SFig 4: Effect of different siASCs on HCV-induced Golgi fragmentation**

Huh7 cells were reverse transfected with two different siRNAs against ASC (siASC) or control siRNA (siCtrl) prior to infection with HCV for 3 and 5 days. (A) Representative images of cells depleted or not of ASC and immunostained with antibodies against GM130 (green) and HCV-Core (not shown). DAPI staining marks nuclei (blue). (B) The characteristics of Golgi fragments were calculated. Data shown are the means ± SD; n=2 independent experiments, >50 cells per condition.

**SFig 5: ASC and GM130 localization in different cell types.**

Representative images of cells non-treated (Ctrl) or treated with Nigericin (0.1 or 1 µM; 3 hours of treatment) and immunostained with antibodies against ASC (green) and GM130 (red). DAPI staining marks nuclei (blue).
